# HDAC3 prevents enhancer hyperactivation to enable developmental transitions

**DOI:** 10.1101/2025.08.09.669452

**Authors:** Nikolaos Stamidis, Anne Wenzel, Konrad Kacper Uściło, Smaragda Kompocholi, Sandra Bages-Arnal, Gemma Noviello, Lea Haarup Gregersen, James Alexander Hackett, Jan Jakub Żylicz

## Abstract

Dynamic gene regulation requires precise cooperation between transcription factors and chromatin modifiers at regulatory elements to achieve not only activation or repression, but also appropriate transcript dosage. However, the molecular mechanisms that ensure developmental genes are transcribed at specific levels remain largely unknown. Here, we discover that the epigenetic repressor histone deacetylase 3 (HDAC3), together with co-activators, binds a subset of the most active putative enhancer elements. Using a tunable mouse embryonic stem cell degron system, we uncover that HDAC3 directly prevents enhancer—and consequently gene—overactivation, effectively establishing a molecular “speed limit” for many naïve pluripotency and housekeeping genes. Interestingly, we find that both the catalytic and non-catalytic functions of HDAC3 contribute to establishing this physiological transcript dose. Specifically during early development, HDAC3 rather than functioning as a canonical repressor, constrains the activity of highly transcribed genes of the implanting epiblast and ensures timely exit from naïve pluripotency *in vitro*. Altogether, this indicates that a dynamic equilibrium between activators and repressors coexists at highly active enhancer elements in pluripotent stem cells, establishing appropriate transcriptional dosage and rendering them responsive to signaling cues, thereby enabling timely and coordinated developmental progression.

## Introduction

Maintenance of cell identity and differentiation depends on precise control of gene expression. This regulation is orchestrated at both enhancers and promoters, primarily by transcription factors (TFs), which act in concert with co-regulators ^1^. Together, they facilitate the loading of RNA polymerase II (RNAPII), often through modulation of chromatin structure ^2,3^. This is particularly relevant at enhancer elements, which not only establish the spatio-temporal pattern of gene expression but also regulate the transcript dosage through promoting RNAPII initiation^4^. Among the various enzymatic activities of the co-regulators that are active at enhancers, acetylation of proteins, primarily histones, plays a central role in transcriptional activation ^1,5^. Notably, histone acetylation is also one of the most rapidly turned-over chromatin modifications, due to the opposing functions of histone acetyl-transferases and deacetylases ^6–8^. Understanding how these opposing functions modulate transcription repression and dosage is particularly important for the dynamically evolving transcriptional states of the developing embryo.

Based on sequence homology and mode of action, the family of histone deacetylase (HDAC) enzymes can be divided into four classes ^8^. Class I HDACs have emerged as important regulators of cellular identity ^9–22^. Indeed, their absence during embryonic development results in gastrulation failure, and early embryonic lethality ^15,19,22,23^. Class I of HDACs comprise HDAC1, HDAC2, HDAC3 and HDAC8, and are known to act as part of large multiprotein co-repressor complexes^8^. While HDAC1 and HDAC2 participate in many such complexes and can mutually compensate for one another, HDAC3 specifically interacts with the nuclear receptor corepressor 1 (NCOR1 or NCOR) or its paralogue the silencing mediator of retinoid acid (NCOR2 or SMRT) ^24^. Together, they are recruited to chromatin primarily by various TFs, such as ETS or AP-1 family members, and ligand-unbound nuclear receptors, including retinoid acid receptor (RAR), thyroid hormone receptor (TR), and PPARγ ^25^ . NCOR1/2 not only bind HDAC3 but are also essential for its catalytic activity ^24,26–28^. Other confirmed partners of the NCOR-HDAC3 co-repressor complex are GPS2 and TBL1 (and/or its homolog TBLR1) ^29–31^. In multiple cellular contexts, the canonical NCOR1/2-HDAC3 complex is recruited to gene regulatory elements ^32^. Specifically in the case of nuclear receptors, ligand binding leads to displacement of the co-repressor complexes and their replacement by co-activator complexes, thereby activating transcription ^33^. Whether this mode of action of HDAC3 also regulates early development remains unclear.

In somatic cells, HDAC3 is involved in multiple physiological and developmental processes ^34^, encompassing both catalytic-dependent and -independent functions. Intriguingly, while genetic ablation of HDAC3 leads to embryonic lethality by gastrulation, mice carrying mutations in the NCOR1/2 interaction domains survive to adulthood, despite lacking detectable catalytic activity^28^. Conditional knockout of HDAC3 in adult mouse liver disrupts liver homeostasis and circadian regulation, and induces liver hepatosteatosis ^27,35–38^. In this system, HDAC3 is recruited to promoters via the nuclear receptor REV-ERBa in a diurnal manner, leading to reduced H3K9ac and activation of genes related to lipid metabolism ^35,38^. However, HDAC3’s catalytic activity and histone acetylation levels are dispensable for lipid metabolism regulation, as a deacetylase catalytic mutant can rescue the phenotype ^27^. In contrast, in brown adipose tissue, both the recruitment and the catalytic activity of HDAC3 are required for the regulation of lipid metabolism-associated genes ^39,40^. Models of early mouse development have shown that the HDAC3-NCOR1/2 complex is involved in cell identity and differentiation of trophoblast stem cells ^41^, endoderm specification ^42^, and X-chromosome inactivation (XCI) in female mouse embryonic stem cells (ESCs) ^43,44^. Nevertheless, the reasons why HDAC3-deficient embryos die during early development, and the molecular mechanisms by which HDAC3 regulates autosomal genes, remain unclear.

Here, using *Hdac3* knock out mice, tuneable acute HDAC3 degron ESCs, chromatin profiling and nascent transcriptomics, we reveal an unanticipated function for HDAC3. We find that it is recruited to the most active enhancers where it imposes a “speed limit” for their activation. Specifically in pluripotent cells, HDAC3 prevents overactivation of many housekeeping and pluripotency genes. In line with this, HDAC3 is vital for timely exit from naïve pluripotency, efficient response to signalling cues and peri-implantation development. Through tethering and complementation experiments we reveal that HDAC3 has both catalytic-dependent and - independent functions, which cooperate to allow cells to transit through differentiation.

## Results

### Histone deacetylase 3 localizes at putative active enhancer elements

Our recent work uncovered that exit from naïve pluripotency is associated with increased histone acetylation turnover ^45^. We reasoned that enzymes allowing for dynamic acetylation changes are likely important in the context of development and pluripotency. We focused on HDAC3, which is expressed throughout development, mediates XCI, and whose loss leads to post-implantation lethality due to unknown reasons ^19,23^. To discern the molecular function of HDAC3 on autosomes, we first asked where in the genome is HDAC3 localized in the context of pluripotent mouse ESCs. For this, we utilized our published ChIP sequencing data for HDAC3 as well as for H3K27ac, H3K4me1, and H3K4me3 chromatin marks generated from female ESCs growing in 2iLIF conditions ^44^. H3K27ac and H3K4me1 co-enrichment typically demarcates active enhancer elements ^46–48^, while H3K27ac and H3K4me3, along with low H3K4me1 levels, mark active promoters ^47,49–51^. Upon peak calling using MACS3, we identified 11,390 HDAC3, 46,052 H3K27ac, 115,531 H3K4me1, and 39,429 H3K4me3 peaks. When focusing on all HDAC3-bound loci, we found that these regions are also co-enriched for enhancer-associated H3K27ac and H3K4me1 marks (**Fig. 1a**). On the other hand, there was only moderate H3K4me3 enrichment at HDAC3-bound sites (**Fig. 1a**). Based on the overlap between the H3K27ac and H3K4me1 peaks, we identified a set of 47,507 putative active regulatory elements, which include both enhancers and promoters. When compared to these regulatory elements, HDAC3 peaks were enriched in intergenic regions and depleted from promoter elements (**Fig. 1b**). Of the intergenic peaks, the median distance from transcription start site (TSS) is approximately 18.4 kb (ranging from 1002 to 905,093 bp – **Fig. 1c**).

**Figure 1.**
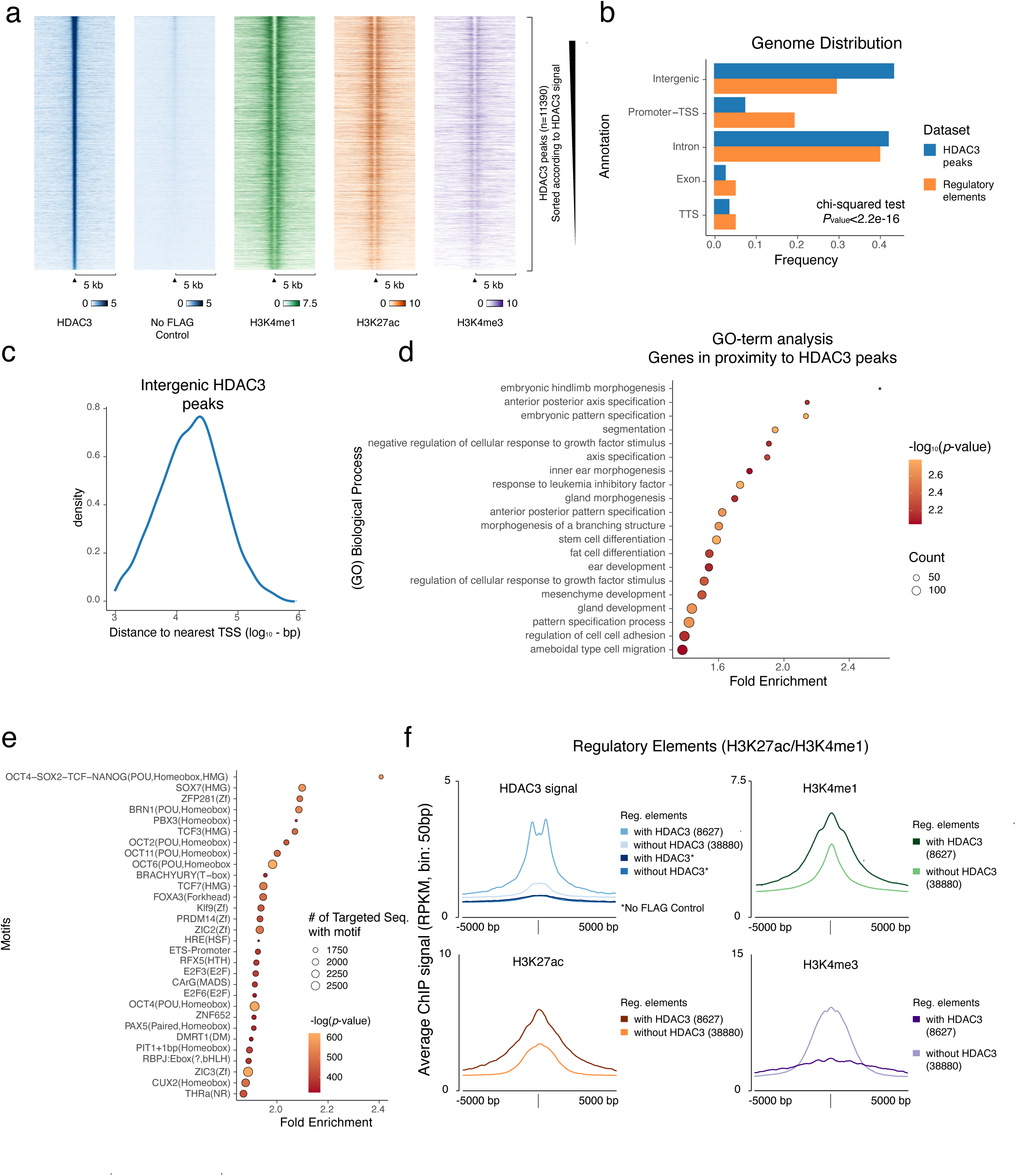
HDAC3 binds to active enhancers. **a**, Heatmaps of all HDAC3 peaks (±5,000 bp) showing enrichment of HDAC3-FLAG and no FLAG control as well as H3K4me1, H3K4me3, and H3K27ac. Dataset was downloaded from Zylicz et al. (GEO GSE116990) ^44^ and originates from female ESCs. Regions are sorted in descending order of HDAC3 signal. Color intensity indicates reads per million per 1 kb (RPKM), binned at 100 bp intervals. **b**, Bar plot illustrating the genomic distribution of HDAC3 peaks and putative regulatory elements. Bar lengths represent percentages. Statistical significance was assessed using the chi-square test. **c**, Density plot showing the distribution of distances from each intergenic HDAC3 peak to the nearest annotated transcription start site (TSS). **d**, Dot plot displaying top 20 enriched Gene Ontology (GO) biological function terms related to genes in proximity to HDAC3 peaks. Dot size indicates the number of genes associated with each GO term, and color intensity represents –log_10_(adjusted *p-*value). P-values were corrected for multiple testing using the Benjamini–Hochberg method. **e**, Dot plot displaying top 30 enriched TF motifs at HDAC3-bound regulatory elements identified using HOMER tool. Dot size indicates the number of regulatory elements with a TF motif, and color intensity represents –ln(*p-*value). **f**, Metaplots of putative regulatory elements with or without HDAC3 binding, showing the average ChIP signal (in RPKM) for HDAC3-FLAG, no FLAG control, H3K4me1, H3K4me3, and H3K27ac, binned at 50 bp intervals and centered on the regulatory element (±5,000 bp).

Further analysis revealed that the genes in proximity to HDAC3 peaks are enriched for Gene Ontology (GO) terms related to both pluripotency (response to leukemia inhibitory factor - LIF), and lineage commitment during gastrulation, i.e. stem cell differentiation, embryonic pattern specification, anterior-posterior pattern specification, segmentation and somite development (**Fig. 1d**). Next, we classified the putative regulatory elements based on the presence or absence of HDAC3 peaks. We found that around 18.2% of the putative active regulatory elements identified in this study contain an HDAC3 peak or localize within 100 bp from one. This comparison further confirmed that HDAC3-bound regulatory regions are predominantly intergenic rather than associated with promoters (**Extended Data Fig. 1c**). This portion of regulatory elements accounts for a potentially highly active set of enhancers, since it is enriched for H3K4me1 and H3K27ac but has lower levels of H3K4me3 (**Fig. 1f**, **Extended Data Fig. 1a, b, d**). Further corroborating our GO term enrichment analysis, we found that HDAC3-bound elements are more likely to contain motifs of core pluripotency-related TFs, when compared to unbound putative regulatory elements (**Fig. 1e**). Altogether, these data imply that in naïve ESCs, HDAC3 predominantly resides in the most active enhancers bound by core pluripotency TFs. This is further corroborated by recent proximity ligation experiments, which profiled proteins residing in proximity to the key histone acetyltransferase P300 in mouse ESCs ^52^. Upon reanalysing this proteomic study, we found that NCOR1, NCOR2 along with HDAC3 and other components of the complex (i.e. TBL1X, TBL1XR1, GPS2) are among the most enriched proteins in proximity to P300 reaching enrichment levels comparable to those of the mediator complex subunits (**Extended Data Fig. 1e**). Overall, we found that in naïve ESCs HDAC3, contrary to its canonical role as a transcriptional repressor, predominantly localizes to highly active autosomal enhancers, which are also co-bound by such co-activators as P300 and the pluripotency TFs.

### HDAC3 regulates highly active regulatory elements and their nearby genes

Having established that HDAC3 co-occupies the most active enhancers with potent transcriptional activators, we set out to assess HDAC3’s role at such sites. To manipulate the levels of HDAC3 in naïve ESCs, we generated two independent clones of an auxin-inducible degron system for HDAC3 in male E14Tg2a ESCs (**Fig. 2a**). These HDAC3-mAID-FLAG ESC lines allow for rapid and acute depletion of HDAC3 within 3 hours of treatment with auxin (indole acetic acid - IAA; **Fig. 2b, Extended Data Fig. 2a**). The use of a male cell line allowed us to focus on the role of HDAC3 in autosomal transcription regulation, and exclude phenotypes related to the XCI. We combined these acute loss-of-function models with transient transcriptome profiling using TT_chem_-seq in order to discern immediate transcriptional effects 3 hours post IAA treatment (**Fig. 2c**) ^53^. Importantly, this method allowed us to study nascent transcription, as well as quantify unstable regulatory element-associated RNAs such as enhancer-associated RNAs (eRNAs) and promoter upstream transcripts (PROMPTs). Such transcripts, produced from regulatory elements, both aid in enhancer element identification and also correlate with their levels of activity ^54–56^.

**Figure 2.**
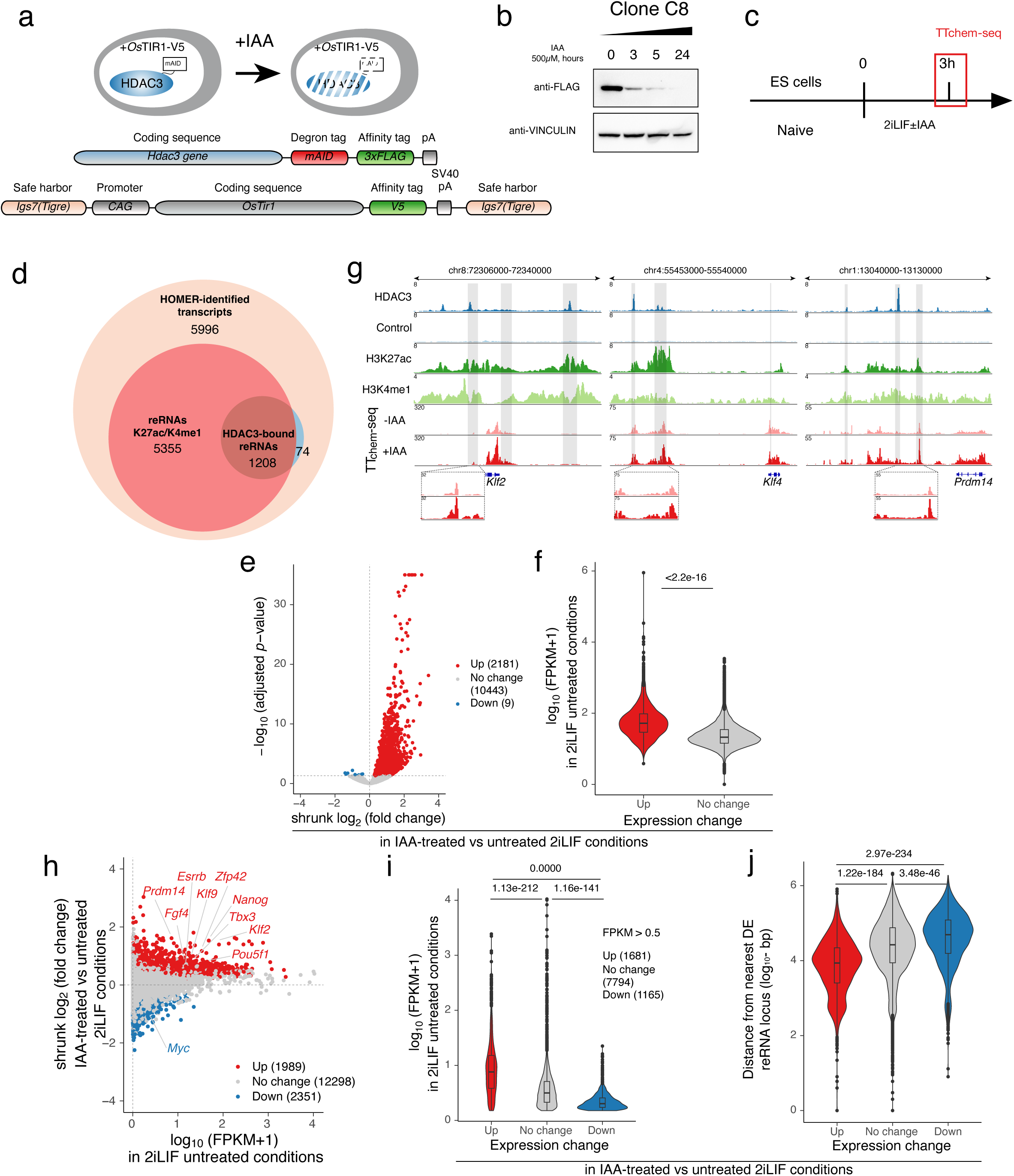
HDAC3 prevents overactivation of highly active enhancers and their nearby genes. **a**, Schematic illustrating the design of the HDAC3 degron system at the endogenous *Hdac3* and *Igs7 (Tigre)* loci. **b,** Immunoblot analysis of whole-cell extracts from HDAC3-mAID-FLAG degron ESCs treated with indole acetic acid (IAA) for varying durations. Blots were probed with anti-FLAG and anti-VINCULIN antibodies. **c**, Schematic of the experimental design for the 3-hour HDAC3 depletion. ESCs cultured in naïve 2iLIF conditions were treated with or without IAA for 3 hours, with a 15-minute 4sU nucleotide pulse prior to harvesting for TT_chem_-seq. **d**, Euler plot showing overlap of HOMER-identified transcript loci with putative H3K27ac/H3K4me1-enriched regulatory elements and HDAC3 peaks. Regulatory element associated RNAs: reRNAs. **e**, Volcano plot of reRNAs in IAA-treated versus untreated 2iLIF conditions with shrunk log_2_(fold change) and -log10(adjusted *p-*value) calculated using DESeq2. Differentially expressed reRNAs with adjusted *p-*value<0.05 are highlighted in color. **f**, Violin plot showing nascent transcription levels in the untreated 2iLIF condition of upregulated reRNAs compared to unchanged reRNAs. Transcript levels are size-factor normalized fragments per kilobase per million reads (FPKMs). Statistical analysis was performed using the Wilcoxon-–Mann–Whitney test. **g,** Genome browser tracks of *Klf2*, *Klf4* and *Prdm14* loci showing HDAC3-FLAG, no FLAG control, H3K4me1 and H3K27ac ChIP signal from published datasets in female ESC ^44^. Shown is also TT_chem_-seq signal in IAA-treated or untreated male HDAC3-mAID-FLAG ESC. In gray, we highlighted enhancer elements identified using STARR-Seq in previous study ^58^. **h**, MA-plot showing the difference in shrunk log2(fold change) of differentially expressed genes in IAA-treated versus untreated 2iLIF conditions against nascent transcription levels in unperturbed 2iLIF degron ESCs. **i**, Violin plot showing nascent transcription levels in the untreated 2iLIF ESCs. Plotted are upregulated, downregulated and unchanged genes upon IAA treatment. Transcript levels are size-factor normalized fragments per kilobase per million reads (FPKMs). Statistical analysis was performed using Kruskal–Wallis rank sum test followed by Dunn’s test for pairwise comparisons. *P-*values were corrected using the Benjamini–Hochberg method. **j**, Violin plot showing the distances from genes to their nearest differentially expressed reRNA, with genes categorized as upregulated, downregulated, or unchanged upon IAA treatment. Statistical analysis was performed using Kruskal–Wallis rank sum test followed by Dunn’s test for pairwise comparisons. *P*-values were corrected using the Benjamini–Hochberg method.

At first, we depleted HDAC3 for 3 hours in self-renewing naïve 2iLIF conditions and used HOMER to identify high confidence nascent transcription from non-genic regions. Specifically in our analysis, we included RNAs that do not overlap with the sense strand of any annotated transcript in ENSEMBLE. To exclude signal from inefficient transcription termination, we also excluded all transcripts mapping to the regions 5kb downstream of all annotated transcription termination sites. We then merged all transcripts identified within 100 bp and selected those that have substantial coverage (mean count > 7, based on DESeq2 independentFiltering), referring to them as HOMER transcripts. Upon overlapping HOMER transcripts with active regulatory elements marked by H3K27ac and H3K4me1 co-enrichment, we found 6563 putative regulatory RNA transcripts (reRNAs). Of these, 1208 originate from elements with an HDAC3 peak (**Fig. 2d**). Notably, when analysing from the perspective of putative regulatory elements, we observed that HDAC3 occupies approximately 32% of those that exhibit significant transcriptional activity (**Extended data Fig. 2b**). All in all, we successfully identified reRNAs transcribed primarily from enhancer elements, many of which are bound by HDAC3.

To determine the effect of HDAC3 depletion on reRNA transcription, we performed differential expression (DE) analysis using DEseq2. We found that HDAC3 depletion results in significant upregulation of 2181 HOMER transcripts, in contrast to only 9 downregulated, thereby confirming the role of HDAC3 as a *bona fide* negative regulator of transcription (**Fig. 2e**). Interestingly, under untreated 2iLIF conditions, HOMER transcripts that become upregulated upon HDAC3 loss already exhibited significantly higher basal transcript levels compared to those that remain unchanged (**Fig. 2f**). Gene Set Enrichment Analysis of DE HOMER transcripts showed significant positive enrichment for HDAC3-bound reRNA loci (**Extended data Fig. 2c**). Indeed, HDAC3-bound reRNA loci were twice as likely (2.2 fold) to become upregulated upon IAA treatment compared to HDAC3-unbound sites (**Supplementary Table 1**). This indicates that HDAC3 likely acts *in cis* as a “speed limit” for highly active regulatory elements, rather than repressing poised or inactive sites.

Based on our finding that HDAC3 prevents enhancer overactivation, we hypothesized that HDAC3 could regulate genes that are highly expressed in 2iLIF conditions. To test this, we measured the levels of nascent transcripts produced from genic regions and performed differential gene expression analysis (DGE). Indeed, HDAC3 depletion caused upregulation of 1989 genes (**Fig. 2h**), many of which are related to pluripotency as well as housekeeping cellular functions such as ribosome biogenesis and translation, metabolism, and RNA processing (**Extended data Fig. 2d**). Indeed, such naïve pluripotency loci as *Klf2, Klf4* and *Prdm14* showed upregulation of transcription at both genes and their previously validated enhancer elements (**Fig. 2g**) ^57–60^. At these well-described loci, HDAC3 is predominantly bound to proximal and distal enhancers, however some binding at promoter elements is also observed. Notably, genes that are upregulated upon HDAC3 loss already exhibit significantly higher transcriptional activity in untreated 2iLIF conditions and reside closer to DE HOMER transcript loci compared to HDAC3-unresponsive genes (**Fig. 2i-j**). In line with the possible direct *in cis* regulation of these genes by HDAC3, we found that upregulated genes are also in close proximity to HDAC3 peaks (**Extended data Fig. 2e**). Unlike for the HOMER transcripts, we also identified a number of downregulated genes upon IAA treatment (2351 - **Fig. 2h**). However, these genes on average are transcribed significantly lower in untreated 2iLIF conditions, compared to the rest of the genes, and reside further away from DE HOMER transcript loci (**Fig. 2i-j**). Moreover, downregulated genes are also more remote from HDAC3 peaks (**Extended data Fig. 2e**). This indicates that in ESCs, HDAC3 is unlikely to function as a *cis*-acting transcriptional activator, rather, the observed changes are either due to indirect effects or HDAC3’s *trans*-acting role. In line with our TT_chem_-seq results, an additional EU-labelling experiment confirmed that HDAC3 does not lead to a global increase in transcription levels (**Extended Data Fig. 2f**). Overall, our results reveal that HDAC3 thresholds the maximum expression of pluripotency and house-keeping genes, thereby establishing appropriate transcript dose.

### The buffering role of HDAC3 relies in part on its catalytic activity and is specific to ESCs

To gain further mechanistic insight into the role of HDAC3 in transcriptional regulation, we set out to complement the phenotype of HDAC3 loss. To this end, in the HDAC3-AID-FLAG degron ESC line, we used a PiggyBac-based vector to ectopically express either wild-type HDAC3 or the catalytically inactive HDAC3-Y298F mutant ^27^. Following a 3-hour IAA treatment, we performed TT_chem_-seq to measure nascent transcription (**Fig. 3a-b**). By focusing on HDAC3-dependent genes identified in previous experiments (**Fig. 2h**), we found that the wild-type HDAC3 can fully complement IAA treatment and prevent overactivation of such genes as *Nanog, Esrrb, Klf2* and *Prdm14* among others (**Fig. 3d-e, Extended Data Fig. 3a**). On the other hand, the catalytically inactive HDAC3-Y298F mutant could partially complement the phenotype of endogenous HDAC3 loss (**Fig. 3d-e**). Notably, unbiased principal component analysis revealed that the overexpression of the catalytically inactive HDAC3-Y298F mutant, but not of the wild-type protein, leads to a transcriptional change (**Fig. 3c, Extended Data Fig. 3b-e**). However these alterations are less pronounced than those observed upon HDAC3 loss and do not affect the pluripotency genes. Despite this confounding effect, our results indicate that HDAC3 prevents gene overactivation through its catalytic activity but with the aid of the catalytic-independent function.

**Figure 3.**
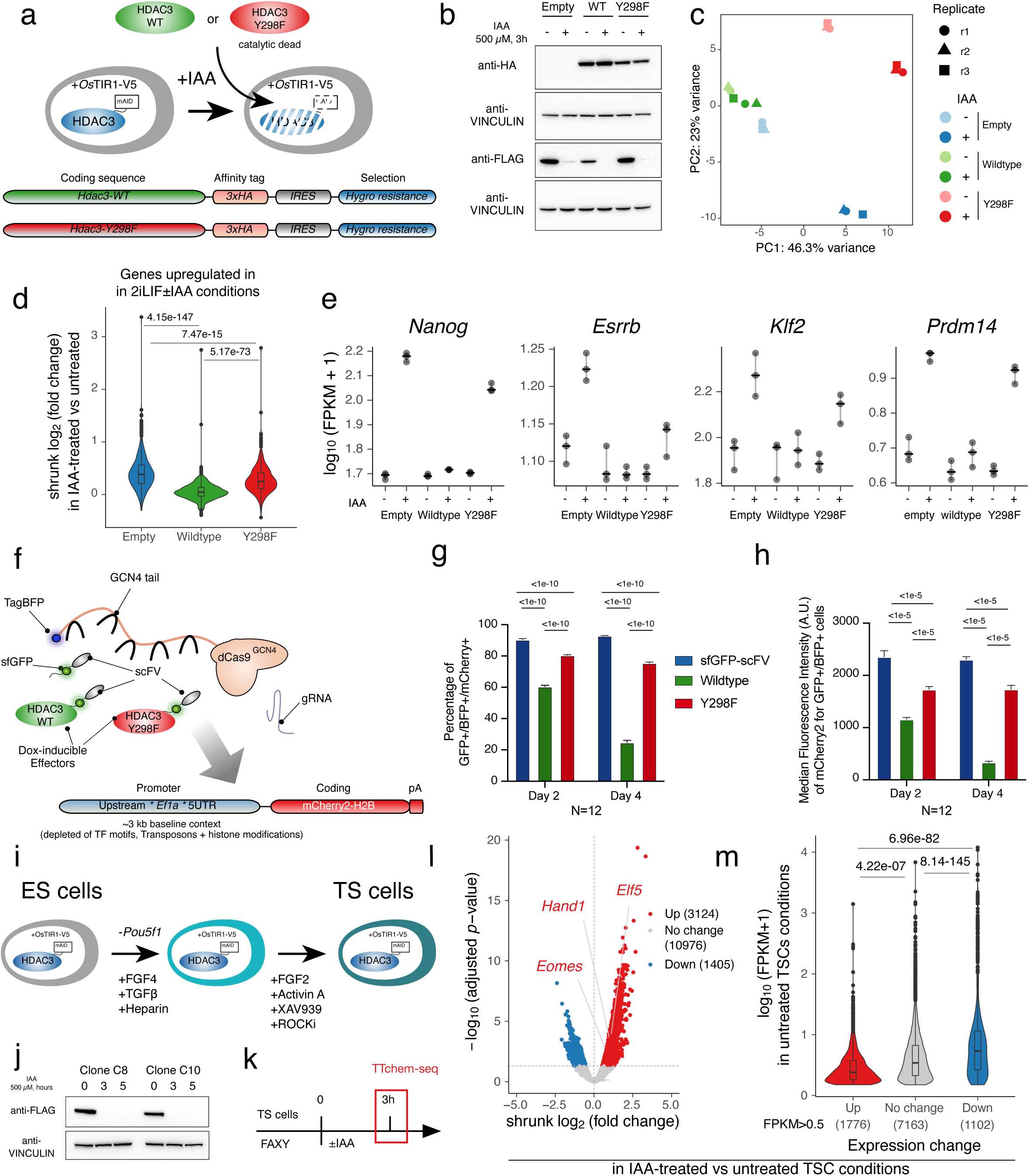
Catalytic activity of HDAC3 prevents gene overactivation in ESCs but not in TSCs. **a**, Schematic of the complementation experiment design, involving transfection with an empty vector, HDAC3^WT^, or the catalytically inactive mutant HDAC3^Y298F^. **b**, Immunoblot analysis of the complementation experiment (**a**) with whole-cell extracts from HDAC3-mAID-FLAG degron ESCs treated with indole acetic acid (IAA) for 3 hours. Blots were probed with anti-FLAG (endogenous HDAC3), anti-HA (transfected HDAC3) and anti-VINCULIN antibodies **c**, Principal Component Analysis (PCA) plot of nascent transcriptomics data from the 3-hour HDAC3 depletion and complementation experiment. **d**, Violin plot illustrating the shrunk log_2_(fold change), in degron ESCs complemented with empty vector, HDAC3^WT^, or HDAC3^Y298F^ upon treatment with or without IAA, for genes that were previously identified as upregulated genes upon HDAC3 loss (Fig. 2h). Statistical analysis was performed using Kruskal–Wallis rank sum test followed by Dunn’s test. *P-*values were corrected using the Benjamini–Hochberg method. **e**, Transcription levels of representative pluripotency-associated genes in ESCs degron line complemented by empty vector, HDAC3^WT^, or HDAC3^Y298F^ and treated with IAA for 3-hours. **f**, Schematic of the dCas9-GCN4-based epigenome editing system. Constitutively expressed dCas9 fused to GCN4-binding sites and TagBFP was stably transfected with doxycycline-inducible vectors expressing sfGFP-scFV, HDAC3^WT^-sfGFP-scFV, or HDAC3^Y298F^-sfGFP-scFV. Guide RNAs were used to target a safe-harbor locus expressing mCherry-H2B under the *Ef1a* promoter. **g–h**, Bar plots showing the percentage of sfGFP+/TagBFP+/mCherry2+ positive cells and their median fluorescence intensity (A.U.) upon doxycycline treatment. Statistical analysis was performed using ANOVA followed by Šidák’s test for multiple comparisons. **i**, Schematic of the protocol for derivation of trophoblast stem cells (TSCs) from ESCs. **j**, Immunoblot analysis of whole-cell extracts from HDAC3-mAID-FLAG degron TSCs treated with indole acetic acid (IAA) for varying durations. Blots were probed with anti-FLAG and anti-VINCULIN antibodies. **k**, Schematic of the experimental design for the 3-hour HDAC3 depletion in TSCs cultured in N2B27 with FGF2, Activin A, XAV939 and Y27632 (FAXY). **l**, Volcano plot of genes in IAA-treated versus untreated TSCs with shrunk log_2_(fold change) and -log10(adjusted *p-*value) calculated using DESeq2. Differentially expressed genes with adjusted p-value<0.05 are highlighted in color. **m**, Violin plot showing nascent transcription levels in the untreated TSCs. Plotted are upregulated, downregulated and unchanged genes upon IAA treatment. Transcript levels are size-factor normalized fragments per kilobase per million reads (FPKMs). Statistical analysis was performed using the Kruskal–Wallis rank sum test followed by Dunn’s test. *P-*values were corrected using the Benjamini–Hochberg method..

Complementation experiments, nascent transcriptomic analysis in combination with HDAC3 profiling, all indicate that HDAC3 acts *in cis* on chromatin to prevent gene overactivation. In order to further test this model, we asked whether *de novo* recruitment of HDAC3 activity is sufficient to regulate gene expression. To this end, we implemented a recently developed ESC line expressing a catalytically inactive dCas9 fused to an array of GCN4 motifs (dCas9-GCN4). This system allows tethering of target proteins, when fused to a single-chain variable fragment (scFV), to a constitutively active promoter driving the expression of mCherry-H2B (**Fig. 3f**) ^61^. As a negative control, we used superfolded GFP (sfGFP) fused to scFV. Additionally, we have established lines which upon doxycycline treatment express full-length wild-type HDAC3 or the HDAC3-Y298F mutant fused to sfGFP and scFV. Upon doxycycline treatment we have observed that wild-type HDAC3, and to a much lesser extent HDAC3-Y298F mutant, is capable of reducing the expression of mCherry-H2B, as assessed both by the percentage of cells in which silencing is observed and the respective fluorescence intensity of the reporter (**Fig. 3g-h**). Together with our complementation assays, these data indicate that *cis*-acting HDAC3, through its catalytic activity, is sufficient to reduce the expression of active genes. Therefore, our experiments reveal that in pluripotent stem cells HDAC3, co-binds with activating complexes (e.g. P300) at the most active regulatory elements to establish a dynamic equilibrium and reduce the transcriptional dose of highly active genes.

Our findings in ESCs are in contrast to previous work performed in other cellular contexts, where HDAC3 maintains gene silencing and its displacement allows for gene activation ^34^. Therefore, we wanted to determine whether the role of HDAC3 in regulating highly transcribed genes is a unique feature of pluripotent stem cells. We decided to focus on the trophoblast stem cells (TSCs), which, similarly to ESCs, approximate early mouse development, but do not express the core pluripotency genes ^62,63^. For this, we *trans-*differentiated HDAC3-AID-FLAG ESCs into TSCs and established two independent clones (**Fig. 3i, Extended Data Fig. 4a**) ^62–66^. Next, we performed TT_chem_-seq upon depletion of HDAC3 for 3 hours (**Fig. 3j-k**). Our DGE analysis showed that HDAC3 loss in TSCs results in upregulation of 3124 genes and downregulation of an additional 1405 genes (**Fig. 3l, Extended Data Fig. 4b**). Unlike ESCs, in TSCs, the genes upregulated upon HDAC3 loss showed significantly lower basal expression in untreated TSCs compared to the HDAC3-independent genes (**Fig. 3m**). Furthermore, trophoblast-specific markers *Hand1*, *Elf5*, and *Eomes* were upregulated but not *Cdx2, Krt8 or Gata3* (**Fig. 3l**). Further data integration revealed limited overlap in the genes differentially expressed in ESCs and TSCs upon HDAC3 loss (**Extended Data Fig. 4d**). More strikingly, even the genes highly expressed in both ESCs and TSCs control conditions did not respond to HDAC3 loss in the same way (**Extended Data Fig. 4e-f**). Therefore, in the context of TSCs, HDAC3 acts similarly to what was described for somatic cells, where it maintains the repression of genes, preventing their activation. All in all, the role of HDAC3 in regulating highly transcribed genes seems specific to pluripotent cells and in part relies on its catalytic activity.

### HDAC3 follows gene expression changes allowing for robust response to signalling cues

Our work up to this point focused on the role of HDAC3 in self-renewing conditions. Next, we set out to address what the role of HDAC3 is when cells undergo cell-state transitions. In particular, we aimed to understand if there is a role for HDAC3 co-binding with transcriptional activators at naïve pluripotency genes when cells respond to signalling cues. For naïve pluripotent stem cells to become competent for differentiation, they need to undergo a process described as capacitation ^67–70^. This is characterized by extensive chromatin and gene expression changes, shutdown of naïve pluripotency genes with maintenance of core pluripotency TFs ^71,72^. By treating 2iLIF ESCs with FGF2 and Activin A (FA), we can induce epiblast-like cells (EpiLCs), which faithfully recapitulate epiblast capacitation of the E5.5-E6.25 ^67,68,70^. Initially we asked what role HDAC3 plays in mediating acute response to signalling cues. To this end, we treated HDAC3-AID-FLAG ESCs with FA for 3 hours in the presence or absence of IAA, and then performed TT_chem_-seq (**Fig. 4a-b**). DGE analysis comparing FA to 2iLIF conditions validated the induction process. Indeed, we found genes that are known to be immediate targets of ERK-signalling, downstream of FGF receptor, to be upregulated such as *Egr1*, *Arc*, *Dusp4/6*, *Plaur* and *Myc* ^73^ (**Extended Data Fig. 5a-b**). Concomitantly, there was downregulation of such naïve pluripotency markers as *Nanog, Klf2/4, Esrrb and Prdm14* (**Extended Data Fig. 5b**). Having validated the induction, we subsequently focused on the phenotypes linked to HDAC3 loss. We identified 4443 upregulated genes and 2809 downregulated genes in FA conditions upon IAA addition (**Fig. 4c**). In line with the results in 2iLIF conditions, HDAC3 loss combined with FA treatment also resulted in upregulation and downregulation, of highly and lowly expressed genes respectively (**Extended Data Fig. 5c**). Moreover, the highly expressed genes that are differentially expressed associate to housekeeping biological processes such as RNA splicing, translation initiation, and ribosome biogenesis (**Extended Data Fig. 5d**). Interestingly, Gene Set Enrichment Analysis (GSEA) revealed that developmental genes downregulated upon FA induction alone, are significantly enriched in genes that become upregulated upon HDAC3 loss in FA treated cells (**Fig. 4d**). Similarly, genes upregulated upon FA induction alone are significantly enriched for genes that are downregulated in combined IAA and FA treatment. Together these data reveal that HDAC3 is crucial to efficiently respond to signalling cues and initiate the exit from naïve pluripotency. Strikingly, when we verified individual naïve pluripotency genes, which are downregulated upon the 3-hour FA induction, we observed that the levels of nascent transcription were even higher than the levels in the starting 2iLIF conditions (**Fig. 4e**). These effects demonstrate that the HDAC3-dependent hyperactivation of highly transcribed genes impacts the response to signalling not only by preventing silencing upon induction but also over-activating genes above their physiological baseline.

**Figure 4.**
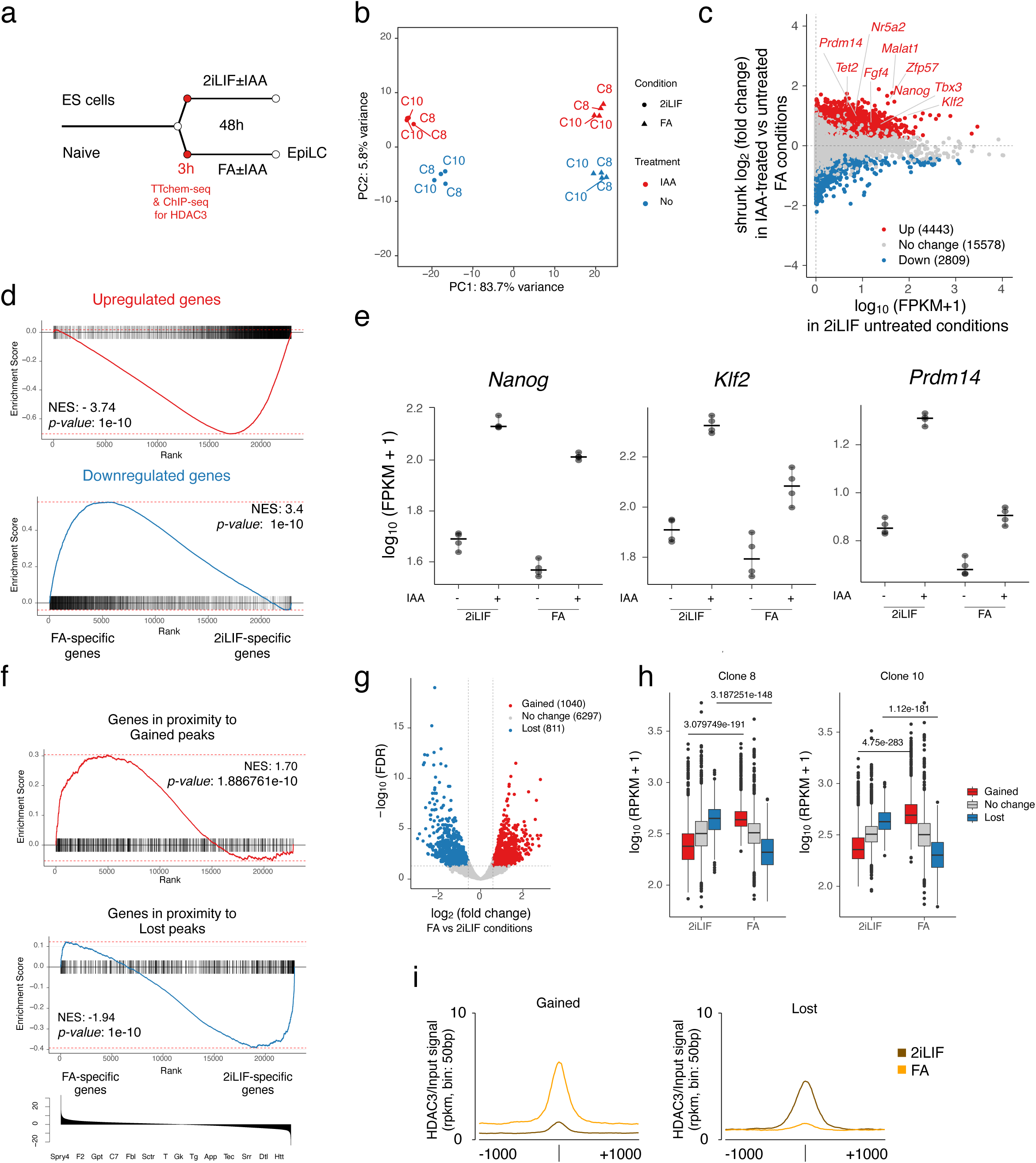
HDAC3 loss prevents exit from naïve pluripotency in mouse embryonic stem cells. **a**, Schematic of the experimental workflow for ESCs exiting naïve pluripotency using EpiLC induction. Naïve ESCs in 2iLIF were switched to FGF2 and Activin A (FA) for 3 hours, with or without IAA, and samples were collected for TT_chem_-seq. Untreated 2iLIF cells and cells treated for 3 hours with FA, were also collected for ChIP-seq analysis. **b**, PCA plot showing gene transcriptomic profiles of different conditions analyzed by TT_chem_-seq. **c**, MA-plot showing shrunk log_2_(fold change) of differentially expressed genes in IAA-treated versus untreated FA conditions against nascent transcription levels in unperturbed 2iLIF degron ESCs. **d**, Gene Set Enrichment Analysis (GSEA) of ranked differentially expressed genes upon FA induction of IAA-untreated ESCs, for genes upregulated or downregulated upon HDAC3 loss in the FA sample. Ranking was based on DESeq2 stat values. **e**, Changes in nascent gene transcription upon HDAC3 loss in 2iLIF±IAA versus FA±IAA for selected naïve pluripotency markers. **f**, GSEA for genes near HDAC3 peaks that gained or lost ChIP signal during the 3-hour FA induction, using DE genes upon FA induction of IAA-untreated ESCs as the reference. Ranking was based on DESeq2 stat values. **g**, Volcano plot illustrating the distribution of HDAC3 ChIP signal changes at HDAC3 peaks during the 3-hour FA treatment. **h**, Box plots showing HDAC3 ChIP signal at peaks that gained, lost, or showed no change in HDAC3 signal during FA treatment. Statistical analysis was performed using Kruskal–Wallis rank sum test followed by Dunn’s test for pairwise comparisons. *P*-values were corrected using the Benjamini–Hochberg method. **i**, Tracks showing average HDAC3 ChIP signal on both strands, binned at 50 bp intervals and centered on gained or lost HDAC3 peaks (±1,000 bp).

Having established that HDAC3 is important for ESCs to respond to signalling cues, we next asked how HDAC3 binding responds to FA treatment. To this end we performed ChIP-seq for HDAC3 in 2iLIF conditions and at 3 hours post FA induction. We used MACS3 to identify peaks in both conditions and then merged them together. Subsequent differential peak enrichment analysis revealed that within the 3-hour FA time-window, HDAC3 signal is partially repositioned (**Fig. 4f-g**). Strikingly, peaks with decreased HDAC3 binding upon FA treatment are preferentially located near to developmental genes becoming downregulated upon 3-hour FA treatment in control conditions (**Fig. 4 f-j**). This implies that HDAC3 becomes rapidly unloaded from regulatory elements upon their silencing. On the other hand, we observed some increased binding of HDAC3 in proximity to genes that are upregulated upon FA treatment alone (**Fig. 4 f-j**). Collectively, we find that HDAC3 is pre-bound on the most active regulatory elements, where it establishes a dynamic equilibrium with transcriptional co-activators. In this context, HDAC3 prevents overactivation of the most active genes, while maintaining their responsiveness to developmentally programmed signalling cues. Additionally, in pluripotent cells HDAC3 binding follows the underlying enhancer activation status, leaving sites upon their silencing and becoming recruited to newly activated regulatory elements.

### Loss of HDAC3 delays exit from naïve pluripotency

While HDAC3 is important for ESCs to acutely respond to FA, it is unclear if this is relevant for functional cell state transitions. To identify the long-term effects of HDAC3 loss during capacitation we performed RNA-seq in HDAC3-AID-FLAG Day 2 EpiLCs induced in the presence or absence of IAA (**Fig. 5a, Extended Data Fig. 6a**). We identified 1015 upregulated genes and 1198 downregulated genes (**Fig. 5b**). Notably, these numbers differ compared to the number DE genes that we identified at the 3-hour time point post FA treatment, implying the existence of potential compensatory mechanisms. Moreover, there is a significant but not complete overlap between the DE genes from the two transcriptomics datasets, most likely due to secondary effects of HDAC3 loss (**Extended Data Fig. 6b, Supplementary Table 3**). Nevertheless, GSEA showed that genes upregulated upon HDAC3 loss are found significantly enriched in genes downregulated upon EpiLC induction alone. The inverse is also true, as genes downregulated upon HDAC3 loss are found enriched in genes upregulated upon EpiLC induction (**Fig. 5d-e**). These data confirmed that ESCs cannot efficiently exit naïve pluripotency in the absence of HDAC3. This phenotype is apparent both in inability to efficiently silence genes specific to naïve pluripotency, and in the failure to establish the EpiLC-specific transcriptional profile (**Fig. 5c-e**). We confirmed our results by immunofluorescence for TBX3, a naive pluripotency marker, which fails to become completely silenced in the absence of HDAC3 (**Extended Data Fig. 6c-d**). Next, we set out to test if this apparent delay in exiting naïve pluripotency leads to functional outcomes. To this end, we performed short reversion experiments, where ESCs are induced towards EpiLCs for 1 day and then returned to 2iLIF conditions for another 24 hours. Without IAA treatment, cells are not able to efficiently re-express TBX3 and form dome-shaped colonies. On the other hand, HDAC3 loss rendered cells more plastic and allowed them to readily upregulate TBX3 in response to 2iLIF and establish characteristic morphology (**Extended Data Fig. 7a-c**). All in all, HDAC3 loss prevents ESCs from efficiently exiting naïve pluripotency at both molecular and functional levels.

**Figure 5.**
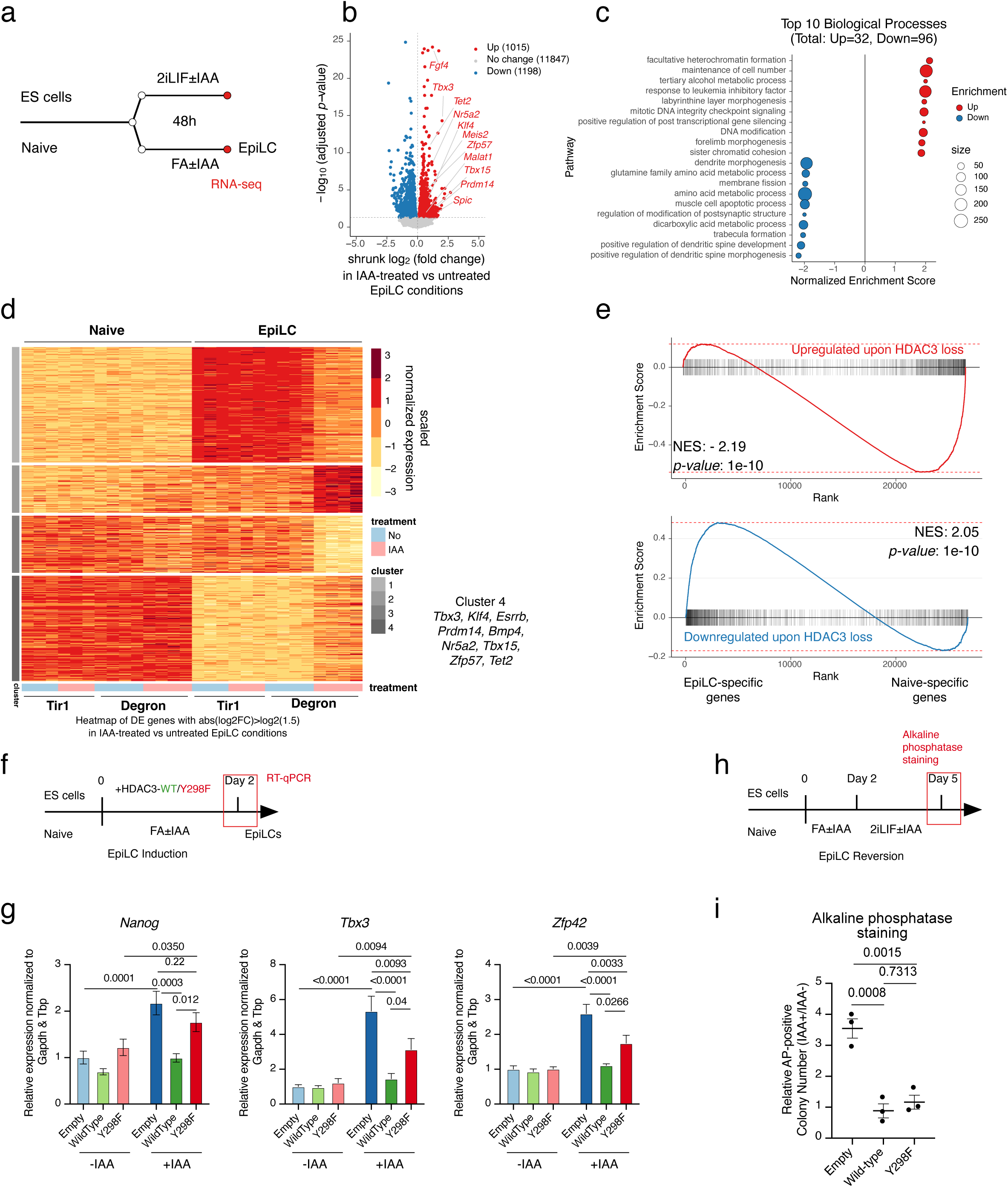
Long term loss of HDAC3 impairs exit from naïve pluripotency and capacitation. **a**, Schematic overview of experimental design. Mouse ESCs were cultured in 2iLIF and induced into EpiLCs with or without IAA. RNA-seq and RT-qPCR analyses were performed 48 hours post-induction. **b**, Volcano plot analysis of gene expression levels in IAA-treated versus untreated EpiLCs with shrunk log_2_(fold change) and -log10(adjusted *p-*value) calculated using DESeq2. Differentially expressed genes with adjusted *p*-value<0.05 are highlighted in color. **c**, GSEA for genes associated with GO terms of biological processes, using ranked DE genes from EpiLC induction with or without IAA. Top enriched pathways are shown. **d**, Heatmap showing scaled normalized expression of differentially expressed genes with absolute shrunk log2(fold change) > log2(1.5) in IAA-treated versus untreated EpiLCs. Cells expressing only TIR1 (not HDAC3-mAID-FLAG) served as controls. **e**, Gene Set Enrichment Analysis (GSEA) of ranked differentially expressed genes upon 2-day induction of IAA-untreated EpiLCs, for genes upregulated or downregulated upon HDAC3 loss in the FA sample. Ranking was based on DESeq2 stat values. **f,** Schematic of experimental workflow as in (a), complemented with vectors encoding HDAC3^WT^, HDAC3^Y298F^, or empty vector. **g**, RT-qPCR analysis of key naïve pluripotency markers in EpiLCs as in (e). Statistical analysis was performed using two-way ANOVA followed by Tukey’s test. **h**, Schematic of extended reversion experiment. Naïve ESCs were induced to EpiLCs for two days, then reverted in 2iLIF for 5 days, and colonies were stained for alkaline phosphatase. **i**, Alkaline phosphatase staining of ESCs after EpiLC induction and reversion, demonstrating the effect of HDAC3 loss on maintenance and re-acquisition of naïve pluripotency. Statistical analysis was performed using one-way ANOVA followed by Tukey’s test.

Under self-renewing conditions wild-type HDAC3, and to a lesser extent the HDAC3-Y298F mutant, can complement the loss of endogenous HDAC3. Next, we set to address, whether the catalytic activity of HDAC3 is needed for efficient EpiLC induction. For this we used the HDAC3-AID-FLAG ESCs complemented with an empty vector or vectors encoding for wild-type HDAC3 or the HDAC3-Y298F mutant. These cells were induced for 2 days toward EpiLC in the presence or absence of IAA. RT-qPCR analysis for the HDAC3-sensitive naïve pluripotency genes revealed that constitutive expression of wild-type HDAC3 complements HDAC3 loss and allows the establishment of silencing of *Nanog, Tbx3* and *Zfp42* (**Fig. 5f-g**). On the other hand, HDAC3-Y298F mutant can partially complement the phenotype of HDAC3 loss, suggesting both catalytic and non-catalytic functions of HDAC3 are at play. Following this, we performed a series of short reversion experiments, as described previously, this time by complementing the loss of endogenous HDAC3 with an empty vector, wild-type HDAC3 or catalytic inactive HDAC3-Y298F (**Extended Data Fig. 6e**). These experiments confirmed that the catalytic function of HDAC3 is primarily, but not solely, required for efficient exit from naïve pluripotency (**Extended Data Fig. 6f**). We further explored this phenotype by performing a set of extended reversion experiments. Briefly, we have induced the complemented ESC lines toward EpiLCs for 2 days in the presence or absence of IAA. This was followed by a reversion in 2iLIF conditions and a colony formation assay with alkaline phosphatase staining as a readout (**Fig. 5h**). While the efficiency of reversion varied across lines in untreated conditions, stable expression of wild-type HDAC3 complemented the IAA phenotype (**Fig. 5i, Extended Data Fig. 6g**). Similarly, the HDAC3-Y298F mutant could largely complement for the loss of endogenous HDAC3 (**Fig. 5i**). Collectively, our results show that HDAC3 is a potent regulator of naïve pluripotency genes and is important for efficient exit from naïve pluripotency. Our complementation experiments found that both catalytically dependent and independent functions regulate naïve pluripotency genes and that HDAC3-Y298F mutant allows for functional exit from naïve pluripotency.

### Genetic ablation of HDAC3 results in embryonic lethality at the time of implantation

Lastly, we asked to what extent our results recapitulate HDAC3’s role in development. To address this question, we generated *Hdac3* maternal and zygotic knockout (KO) embryos. We crossed heterozygous *Hdac3^+/-^*male mice with *Hdac3^loxP/loxP^* females also carrying oocyte-specific *ZP3-cre* (**Fig. 6a**)^74,75^. This strategy allows for the generation of *Hdac3* KO and heterozygous (Het) embryos devoid of maternal protein contribution, which could potentially mask early embryonic phenotypes. Previous studies have reported that genetic ablation of HDAC3 results in embryonic lethality by E9.5 ^19,23^. To characterize the molecular events that precede lethality, we studied the effects of HDAC3 loss at the pre-implantation expanded blastocyst stage of the embryonic (E) day 4.5 as well as at the onset of gastrulation at E6.5.

**Figure 6.**
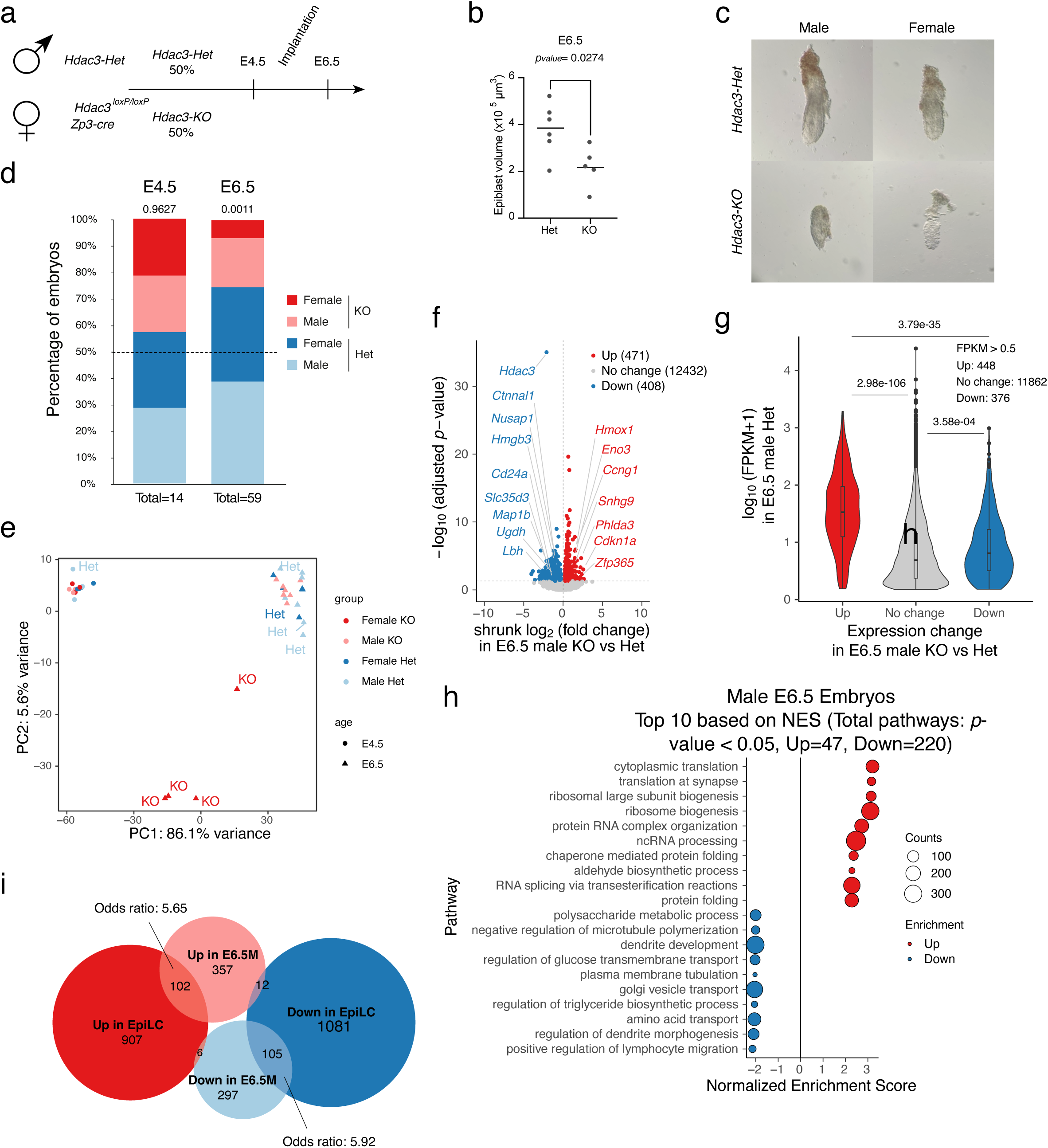
HDAC3 is essential for early mouse embryonic development. **a**, Schematic of breeding strategy and genotypes used to generate *Hdac3* heterozygous (Het) and knockout (KO) embryos, with Zp3-cre for conditional deletion in oocytes. Specifically, male *Hdac3^+/-^* mice were mated with female *Hdac3^loxP/loxP^ R26:Zp3Cre+ve* mice, and embryos were collected at E4.5 and at E6.5 stages. **b**, Quantification of epiblast volumes at E6.5 for *Hdac3-*Het and *Hdac3-*KO embryos. Line indicates the median value and statistical analysis was performed using unpaired t-test. **c**, Representative images of dissected *Hdac3-*Het and *Hdac3-*KO embryos (male and female) by brightfield microscopy. **d**, Genotype distribution of embryos at E4.5 and E6.5, showing expected Mendelian ratios at E4.5 and reduced KO embryos at E6.5 for both sexes. *P-*values were calculated using Chi-squared test. **e**, PCA plot of transcriptomic profiles from male and female *Hdac3-*Het and *Hdac3-*KO embryos at E4.5 and E6.5. **f**, Volcano plot analysis of gene expression levels in E6.5 male *Hdac3-*KO vs. *Hdac3-*Het epiblasts with shrunk log_2_(fold change) and -log10(adjusted *p-*value) calculated using DESeq2. Differentially expressed genes with adjusted p-value<0.05 are highlighted in color. **g**, Violin plot showing gene expression levels in the *Hdac3-*Het male E6.5 epiblasts. Plotted are upregulated, downregulated and unchanged genes in *Hdac3-*KO vs. *Hdac3-*Het E6.5 male epiblasts. Transcript levels are size-factor normalized fragments per kilobase per million reads (FPKMs). Statistical analysis was performed using the Kruskal–Wallis rank sum test followed by Dunn’s test. *P-*values were corrected using the Benjamini–Hochberg method. **h**, GSEA of differentially expressed genes in E6.5 *Hdac3-*KO vs. *Hdac3-*Het male epiblasts for genes associated with GO terms of biological processes. **i**, Euler plots showing overlap of up- and downregulated genes in *Hdac3-*KO male E6.5 epiblasts and in epiblast-like cells (EpiLCs), with calculated odds ratios.

When focusing on E4.5 stage, we observed normal morphology and expected Mendelian ratios (**Fig. 6d**). On the other hand, by E6.5, we found a partially penetrant loss of *Hdac3-KO* embryos with multiple empty deciduae detected (**Fig. 6d**). This wave of embryo loss affects both sexes, but females are more severely affected, as assessed by the deviation from the Mendelian ratios (**Fig. 6d**). Indeed, we estimate an implantation failure of approximately 81% in female and 52% in male *Hdac3-KO* embryos, respectively. Of the rare embryos that escape this implantation defect, *Hdac3-K*O E6.5 embryos are significantly smaller, including in terms of epiblast size (**Fig. 6b-c**, **Extended Data Fig. 8a-b**). Interestingly, despite having smaller epiblasts, surviving *Hdac3-KO* embryos can still sequester NANOG expression at the posterior part of the epiblast, which implies that they are capable of anterior-posterior axis patterning (**Extended Data Fig. 8a**) ^76–78^. Overall, we find that while the maternal pool of HDAC3 is dispensable for early development, zygotically encoded HDAC3 is vital for efficient implantation and post-implantation embryo growth, with female embryos being more severely affected than males.

To better understand the role of HDAC3 during implantation, we performed full-length transcriptome analysis using FLASH-seq ^79^, on whole E4.5 blastocysts and dissected epiblasts at E6.5. Our results found that no major transcriptional changes occur by E4.5, as assessed by PCA and DGE analysis (**Fig. 6e, Extended Data Fig. 8h-i**). On the other hand, at the E6.5 stage, female *Hdac3-KO* epiblasts exhibit dramatic differences in transcript levels compared to *Hdac3-Het* (**Fig. 6e, Extended Data Fig. 8c-d**). This striking phenotype in female embryos is likely a composite effect of defects in XCI as well as an altered autosomal gene expression program. To focus on autosomal gene regulation, we next analyzed male E6.5 epiblasts. DGE analysis identified 471 upregulated genes and 408 downregulated genes (**Fig. 6f**). Among the downregulated genes was *Hdac3*, supporting the robustness of our approach. Our *in vitro* work revealed that in pluripotent stem cells, HDAC3 limits the expression of the most actively transcribed genes. Strikingly, we observed that genes upregulated in male E6.5 *Hdac3-KO* epiblasts compared to *Hdac3-Het* had higher baseline transcript levels than HDAC3-independent genes (**Fig. 6g**). This finding indicates that, also *in vivo*, HDAC3 prevents gene overactivation. Additionally, GSEA for GO terms of biological processes showed that upregulated genes in male E6.5 epiblasts are enriched in genes related to housekeeping functions, similar to what we observed *in vitro* (**Fig. 6h**). Furthermore, despite stark experimental differences, there is significant overlap between not only E6.5 DE genes and the RNA-sequencing results from Day 2 EpiLCs, but also with our nascent transcriptomics datasets (**Fig. 6i, Extended Data Fig. 8f-g, Supplementary Table 3**). All in all, HDAC3 loss results in embryonic lethality at a stage earlier than expected—around the time of implantation. In addition, while we observe a more severe effect in female embryos, HDAC3 loss also affects male embryos in terms of survival, morphology, and gene expression. Finally, the phenotype observed in surviving E6.5 embryos particularly affects highly abundant transcripts related to housekeeping biological processes, in accordance with results from our *in vitro* models.

## Discussion

Gene regulation in all cellular contexts extends far beyond simple binary on/off mechanisms. Instead, precise control of transcriptional dosage underpins the establishment of gene regulatory networks. This fine-tuned regulation enables the formation of attractor cellular states that define cellular identity. At the same time, these states remain inherently poised for change—partially destabilized to allow cells to respond dynamically to external cues, particularly during developmental transitions. Transcriptional dosage is crucial and therefore is established at multiple, parallel levels, including through TFs’ cooperativity and chromatin modifiers. In this study, we find that transcriptional dosage is fine-tuned by epigenetic repressors. Indeed, we reveal that co-binding of transcriptional co-activators such as P300 and co-repressors like HDAC3 to the most active subset of enhancer elements is a key mechanism for achieving appropriate transcriptional output. Our *in vivo* and *in vitro* loss-of-function experiments support a model where HDAC3 acts as a molecular “speed limit”, constraining the expression of highly active pluripotency and housekeeping genes. This inherent destabilization of transcriptional output at the most active genes renders them responsive to signalling cues, enabling timely and coordinated developmental progression.

In line with this model, we found that in naïve ESCs, HDAC3 is bound to approximately 18% of putative active regulatory elements, primarily at non-promoter regions. Indeed, we previously reported HDAC3 binding to regulatory elements on the X chromosome and proposed that HDAC3 is pre-bound in an inactive form, awaiting repressor cues such as Xist RNA binding ^44^. However, these HDAC3-bound loci are also present on autosomes and likely represent a subset of highly active enhancer elements, as indicated by strong enrichment for H3K27ac and H3K4me1, along with a concomitant depletion of the promoter-specific H3K4me3 mark compared to non-HDAC3-bound elements. Apart from the few HDAC3-bound promoters, the remaining H3K4me3 marking is consistent with prior observations that highly active enhancer elements can also be partially decorated by this modification ^51^. The majority of HDAC3-bound putative regulatory elements are located relatively close to the TSS, i.e., 50% of them are within a 19 kb window, and are enriched for binding motifs of key pluripotency TFs. Moreover, genes located near HDAC3-bound regulatory elements are associated with pluripotency, housekeeping functions, and pre-gastrulation processes such as embryonic patterning. Supporting our findings, recent proximity labelling experiments identified NCOR1/2 and HDAC3 as being highly enriched near the co-activator P300 in ESCs ^52^. Furthermore, during rapid adipocyte differentiation, following super-enhancer activation, there is recruitment not only of transcriptional co-activators but also of co-repressors ^80^. Indeed, in brown adipose tissue, HDAC3 is recruited to putative regulatory elements. However, these sites do not correspond to the most active enhancers as indicated by H3K27ac ^81^. In such cellular context, HDAC3 recruitment to regulatory elements can lead to their upregulation, reflecting a non-canonical function. By contrast, in the liver, HDAC3 is in part recruited through ligand-free nuclear receptors to maintain gene silencing ^35,38^. Thus, specifically in ESCs, HDAC3 is bound to the most active putative enhancers to regulate their activity.

HDAC3 binding to the most active putative enhancers raises the question of whether it plays a role under steady state conditions or remains inactive, awaiting signaling cues to initiate gene repression. By leveraging HDAC3 degron ESC lines, nascent transcriptomics, and the observation that a large portion of regulatory elements are transcribed—and that their transcriptional activity reflects their regulatory function ^54–56^—we were able to test these hypotheses. Our integrated data establish HDAC3 as a negative regulator of transcriptional activity in steady-state pluripotent stem cells, where HDAC3 prevents overactivation of reRNAs and the genes located nearby. Indeed, under these steady-state conditions, HDAC3 is functional and helps establish the transcriptional dosage of many highly active genes. These results contrast with previous findings in the liver, where the HDAC3–NCOR1/2 complex functions as a canonical repressor maintaining gene silencing ^35,38^. Indeed, this canonical mode of action appears to dominate in the trophoblast lineage, as HDAC3 loss leads to the upregulation of initially silent genes. This aligns with a recent report showing that NCOR1/2 is recruited by FGF-dependent factor ERF to decommission regulatory elements in TSCs ^41^.

Additionally, the role of HDAC3 in ESCs is distinct from that reported for other class I HDACs, namely HDAC1 and HDAC2. Specifically, rapid loss of HDAC1 in the context of *Hdac2-KO* ESCs leads, among other phenotypes, to the downregulation of highly expressed naïve pluripotency genes ^11^. In line with this observation, short-term treatment of ESCs with trichostatin A (TSA), a potent HDAC inhibitor, results in loss of pluripotency and significant 3D chromatin rearrangement ^82^. Moreover, TSA treatment in breast cancer cell lines also led to drastic downregulation of reRNAs ^83^. In light of these findings, we propose that HDAC3 plays a role that opposes the function of HDAC1/2 by inherently destabilizing naïve pluripotency.

The cellular phenotypes observed upon HDAC3 loss could be linked to its chromatin binding, functioning *in cis* at proximal enhancers. Alternatively, *trans-*acting functions of HDAC3 may also contribute to the observed effects. Two lines of evidence suggest that HDAC3 primarily acts *in cis* to reduce the dosage of naïve pluripotency and housekeeping genes. First, we find that loci encoding reRNAs upregulated upon HDAC3 loss are also enriched for HDAC3 binding, supporting a *cis-*regulatory role. Second, our dCas9 experiments demonstrate that tethering HDAC3 *in cis* to a regulatory element is sufficient to inhibit transcription. Surprisingly, our nascent transcriptomic studies also identified a number of genes that are downregulated upon HDAC3 loss. However, these genes are not located near HDAC3-regulated reRNA loci or HDAC3 peaks and are expressed at significantly lower levels than other genes in unperturbed cells. This is contrary to other systems where HDAC3 is a *cis-*acting transcriptional activator ^81^. Our data could therefore indicate a potential trans-acting role for HDAC3, such as regulating the acetylation status of TFs. Indeed, SIRT1, another deacetylase, has been shown to modulate the activity of the key pluripotency TF SOX2 ^84^ Alternatively, the downregulation of lowly transcribed genes upon HDAC3 loss might result from hyperactivation of already highly active genes. Given the short timepoints used and the absence of a global increase in transcription (as assessed by 4sU-staining), we speculate that hyperactivation of active genes leads to a redistribution of limited transcriptional resources, thereby reducing pervasive transcription of lowly expressed genes.

Another open question is the extent to which HDAC3 regulates gene expression through its enzymatic activity. Our complementation assays, combined with dCas9 tethering experiments, suggest that both catalytic and non-catalytic functions of HDAC3 are important. Indeed, the well-characterized HDAC3^Y298F^ mutant could partially complement the loss of endogenous HDAC3 under steady-state conditions, and to a greater extent during the exit from naïve pluripotency. In line with this finding, tethering the HDAC3^Y298F^ mutant to a reporter gene in ESCs led to modest transcriptional attenuation, unlike wild-type HDAC3. Consistent with these observations, in *Drosophila melanogaster*, enzymatically inactive HDAC3 can partially rescue the developmental phenotype caused by HDAC3 loss ^85^. Similarly, in the mouse liver, loss of endogenous HDAC3 can be partially complemented by the HDAC3^Y298F^ mutant ^27^. These non-catalytic functions of HDAC3 are likely mediated through protein–protein interactions.

Overall, our nascent transcriptomic datasets uncovered HDAC3 as a potent thresholding mechanism for the most active putative enhancers in pluripotent stem cells. Differentiation and *in vivo* experiments further confirmed that this function is crucial for cell state transitions. Indeed, genetic ablation of HDAC3 during mouse embryogenesis leads to acute implantation defects, affecting both sexes but with a more severe impact on females. These data imply an important role of HDAC3 in extraembryonic lineages ^41,42^. Remarkably, this phenotype closely mirrors our *in vitro* findings in terms of timing, as the implanting epiblast corresponds to ESCs exiting naïve pluripotency. In surviving male embryos, we observed upregulation of highly transcribed genes and a significant overlap in differentially expressed genes between the *in vivo* and *in vitro* systems. These knockout embryos also exhibit growth defects, including reduced size and epiblast volume, but retain the ability to form the anterior–posterior axis. These findings are broadly consistent with previously reported *Hdac3* knockout lethality prior to E9.5 ^19,23^. The observed sex differences are likely due to the inability of female embryos to efficiently silence the X chromosome, a process that occurs at implantation ^43,72^. Interestingly, the female embryo phenotype is more severe than that observed in *Xist* knockouts, which successfully implant but show developmental delay by E6.5 ^86^. This suggests that the female phenotype results from a combination of autosomal and XCI deregulation.

Our study provides new insights into how chromatin-modifying enzymes influence the execution of transcriptional programs in a context-specific manner. We underscore the importance of enhancer utilization in these processes and demonstrate how HDAC3 can serve as a rheostat for specific gene sets. Future work should elucidate how chromatin-modifying enzymes are recruited to different genomic regions and how differences between cellular contexts contribute to the diversity of transcriptional outcomes.

## Methods

### Cell culture and cell line generation

E14Tg2a (129/Ola, XY) wild-type mouse ESCs, kindly offered by the lab of Joshua Brickman, were used for all experiments, apart from CRISPR-dCas9^GCN4^-based tethering experiments, in which we used the reporter mCherry2-H2B ES cell line (129/B6, XY) developed by Policarpi et al. ^61^. E14Tg2a cells were grown on gelatin, cultured in Serum/LIF conditions, specifically GMEM (ThermoFisher) supplemented with 10% FBS (Merck), 1,000 U mL^−1^ LIF (in-house production), 0.1 mM 2-mercaptoethanol, 0.1 mM non-essential amino acids, 2mM L-alanyl-L-glutamine equivalent (GlutaMax, ThermoFisher), 1 mM sodium pyruvate. For transition to naïve conditions, ESCs were cultured for 12 days on fibronectin-coated plates (Sigma Aldrich), in N2B27 basal culture medium: DMEM/F12 medium (ThermoFisher) and Neurobasal medium (ThermoFisher) 1:1, with the addition of N2 supplement 1/200 (ThermoFisher), B-27 Serum-Free Supplement 1/100 (ThermoFisher), 2mM L-alanyl-L-glutamine (GlutaMax, ThermoFisher), 0.1 mM 2-mercaptoethanol (Merck), supplemented with 1 µM PD0325901, 3 μM CHIR99021, 1,000 U mL^−1^ LIF (in-house production), as described by Mulas et al. ^87^. After transition, cells were maintained and regularly passaged in these conditions. For mCherry2-H2B reporter line, cells were cultured on gelatin-coated plates in N2B27 basal culture medium (NDiff by Takara, y40002), supplemented with 1 µM PD0325901, 3 μM CHIR99021, 1,000 U mL^−1^ LIF (in-house production), 1% FBS (Millipore) and 1% penicillin-streptomycin (ThermoFisher).

For generation of transgenic cell lines, ESCs growing in Serum/LIF were electroporated using the P3 Primary Cell 4D-Nucleofector™ X Kit (Amaxa - Lonza) according to manufacturer’s instructions. For experiments using piggyBAC system cells were transfected using Lipofectamine 2000 according to manufacturer’s instructions for ES cells.

### ESC Differentiation

#### EpiLC induction

EpiLC inductions was performed according to Hayashi et al. ^68^. Briefly, naïve ESCs were washed 3 times in PBS and then passaged into N2B27 medium supplemented with bFGF (12 ng/mL) and Activin A (20 ng/mL) and 1% KSR for 2 days. Medium was changed daily.

#### TSC transdifferentiation

Naïve ESCs growing on fibronectin-coated plates were lipofected with esiRNAs for Pou5f1 (400ngr per 2LxL10L cells), using Lipofectamine 2000 in Optimem. Transfected cells were incubated for 48 h without media change. Cells were then passaged into TX medium (DMEM/F12 without HEPES and L-glutamine, supplemented with TGF-ß1 (2 ng/mL), FGF4 (25 ng/mL), 1 mg/mL Heparin) on laminin-coated plates and maintained until confluency with regular media changes. Emerging iTSC colonies were transferred to FAXY medium (Fgf2 - 12 ng/mL, Activin A - 20ng/mL, XAV939 - 10µM, Y27632 - 5µM) on fibronectin and maintained in the same conditions.

#### Alkaline phosphatase

Alkaline phosphatase staining was performed as described in Mulas et al. ^87^. Briefly cells were converted into EpiLCs, detached with Accutase (Stem Cell Technologies) for 1 min at 37°C, counted and seeded at low density into laminin-coated plates in 2iLIF conditions. Medium was changed every two days and cells were stained after 5 days using the Alkaline Phosphatase Detection Kit for leukocytes (Merck) according to manufacturer’s instructions. Plates were stored O/N at 4oC, in the dark and colonies were counted the day after.

### Immunofluorescence

#### In embryos

Embryos were fixed in 4% PFA with 0.001% PVA for 20 min at 37 °C, then washed 3 times in PBS-T (0.1% Tween-20, 0.001% PVA) for 10 min at room temperature (RT). Permeabilization was performed for 10–15 min in 0.1 M glycine, 0.3% Triton X-100, and 0.001% PVA, followed by three additional PBS-T washes. Embryos were blocked for 4–5Lh at RT in blocking buffer (1% BSA, 0.1% Tween-20, 10% normal serum, 0.001% PVA in PBS), then incubated with primary antibodies (in blocking buffer) for 1–2 days at 4 °C, rocking. After washing (3 times, 10 min each, PBS-T - 0.001% PVA), embryos were incubated with secondary antibodies overnight at 4 °C in the dark. Embryos were washed (3 times, 10Lmin each, PBS-T - 0.001% PVA - DAPI was added during the second wash step at a final concentration of 1µg/ml), and mounted on poly-lysine–coated glass-bottom plates in Vectashield® Antifade Mounting Medium for imaging (Vector Laboratories). Imaging was performed at a Leica Stellaris Microscope.

#### In cells

Cells were seeded in fibronectin-coated Ibidi mini-dishes and cultured in them. The following day, cells were fixed in 4% PFA for 10 min at RT, permeabilized with 0.25–0.5% Triton X-100 for 10 min, and blocked for ≥3 h at RT (1% BSA, 0.1% Tween-20, 10% normal serum in PBS). Primary antibodies were applied in blocking buffer and incubated overnight at 4L°C. After three PBS-T washes, secondary antibodies were applied for 3 h at RT. Cells were washed again (DAPI added in the second wash) and mounted in Vectashield®. Imaging was performed at a Leica Stellaris Microscope.

### Western Blotting

#### Protein extraction

Cells were detached in 1 mL TryPLE, neutralized in equal volume medium and washed twice in PBS and snap-frozen as dry pellets in dry ice. Cell pellets were lysed in RIPA buffer (1x PBS, 150 mM NaCl, 1% NP40, 0.1% SDS, 0.5% Sodium Deoxycholate, 50 mM Tris-HCl, pH 8.0) supplemented with proteinase inhibitors (cOmplete^TM^, Roche), incubated for 15 minutes on ice, and sonicated using a Bioruptor Plus (Diagenode), set to high power for 10 cycles (30 s ON, 30 s OFF). Protein quantification was performed using Pierce BCA kit (ThermoFisher).

#### SDS-PAGE and immunoblotting

Protein concentration was equalized across samples and 20µg were loaded on a NuPAGE 4-12% Bis-Tris (ThermoFisher) pre-cast gel, for ∼180 minutes at 120-140V, in NuPAGE MED SDS running buffer. Proteins were transferred into a nitrocellulose mebrane 0.45 µm (Amersham^TM^ Protran^®^) for 70 min at 350 mA, blocked for 1 hour with 5% milk in PBS + 0.3% NP40. Primary antibodies were incubated O/N in 1% milk in PBS + 0.3% NP40 at 4°C at indicated concentrations (**Supplementary Table 2**). Membranes were washed 3 times in PBS + 0.3% NP40, incubated for an hour with HRP-conjugated secondary antibody (1:10.000 in 1% milk in PBS + 0.3% NP40).

### Real Time RT-qPCR

#### RNA isolation and RT–qPCR

Total RNA was extracted using RNAeasy Plus Mini kit with DNA digestion and equal amounts of RNA were retrotranscribed into cDNA using Superscript III Reverse Transcriptase. Real Time RT–qPCR was performed using the Power Up Master Mix (Applied Biosystems) in a Light Cycler thermocycler (Roche). Primer efficiency was assessed by standard curves of serially diluted, purified amplicon fragment. All samples were run in quadruplicates; data were analyzed using the 2^-ΔΔCt^ method and normalized to the geometric mean of the signal of housekeeping genes i.e. *Tbp* and *Gapdh*.

### ChIP sequencing

#### ChIP sequencing

Analysis of chromatin occupancy for HDAC3 was performed as described in Zylicz et al. ^44^. Briefly, naïve ESCs were grown to ∼80% confluency, washed 3 times in PBS and switched to N2B27 medium supplemented with Fgf2 (12 ng/mL), Activin A (20ng/mL) and KSR 1% for 3 hours. Cells were incubated for 30 min at room temperature (RT) in 1.5LmM ethylene glycol bis(succinimidyl succinate) (EGS; Thermo Fisher) freshly diluted in PBS and then crosslinked with freshly opened formaldehyde for 10 min at a final concentration of 1.88%. Crosslinking was quenched with 0.11 M Glycine for 5 min at RT. Cells were washed twice in ice-cold PBS, scraped, pelleted by centrifugation (500 xg, 5 min, 4 °C), and snap-frozen in liquid nitrogen.

Frozen pellets were thawed on ice and lysed in Nuclear Lysis Buffer (5 mM MgCl₂, 10LmM Tris-HCl pH 8.0, 1 mM EDTA, 0.1 M sucrose, 0.5% Triton X-100, supplemented with protease inhibitors – cOmplete^TM^, Roche) for 10 min on ice. Cells were homogenized using a dounce homogenizer with 20 strokes each of pestle A and B to release nuclei, which were then pelleted (500g, 15 min, 4 °C) and resuspended in Chromatin Lysis Buffer (150 mM NaCl, 10LmM Tris-HCl pH 8.0, 1 mM EDTA, 0.5 mM EGTA, 0.5% N-lauroylsarcosine, with protease inhibitors).

Chromatin was sheared using a Bioruptor Plus (Diagenode), set to high power for 12 cycles (30 s ON, 30 s OFF) to obtain fragments of ∼300–400 bp. Lysates were centrifuged (500g, 15 min), and the supernatant was diluted in Dilution Buffer (150 mM NaCl, 10LmM Tris-HCl pH 8.0, 6 mM EDTA, 0.125% sodium deoxycholate, 1.25% Triton X-100, with protease inhibitors). A small aliquot was reserved as input control, and the remainder was incubated overnight at 4 °C with 10 μg of anti-Flag M2 antibody and protein G Dynabeads (Invitrogen) pre-washed in buffer.

Immunocomplexes were washed five times (10 min, 4 °C, rotation) in each of two wash buffers: LiCl Wash Buffer (0.5 M lithium chloride, 10 mM Tris-HCl pH 8.1, 1 mM EDTA, 1.1% sodiumdeoxycholate, 1% NP-40), High Salt Wash Buffer (1 M NaCl, 20 mM Tris-HCl pH 8.1, 2 mM EDTA, 0.1% sodium deoxycholate,0.1% SDS, 1% Triton X-100). Final washes were performed in TE-low EDTA buffer (10 mM Tris-HCl pH 8.0, 0.1 mM EDTA), and DNA was recovered in Proteinase K buffer (20 mM HEPES pH 7.9, 1 mM EDTA, 0.5% SDS, 0.8Lmg/mL Proteinase K) for 30 min at 56 °C. Samples were further digested by addition of Proteinase K and NaCl (final concentration 0.3 M NaCl) for 2 h at 56 °C, followed by 4Lh at 68L°C.

#### DNA purification

DNA was purified using paramagnetic beads. Equal volumes of Ampure XP beads and PEG/NaCl solution (final: 13% PEG8000, 0.75 M NaCl) were added to the ChIP reaction, incubated for 15 min at RT, and beads were separated using a magnetic rack. Beads were washed twice with 80% ethanol, air-dried, and DNA was eluted in TE-low EDTA buffer (10 mM Tris-HCl pH 8.0, 0.1 mM EDTA). DNA concentration was measured using Qubit dsDNA HS Assay (Invitrogen) and fragment size assessed using a TapeStation D5000 assay (Agilent).

Libraries were performed using the Ovation® Ultralow System V2 1-16 library preparation kit by NuGEN and single-end sequencing was performed P2-100bp NextSeq2000 kit.

#### Data processing

Raw sequencing data were demultiplexed and converted into FASTQ files using Illumina bcl2fastq Conversion Software (v2.20.0.422). Resulting FASTQ files were processed using nf-core/chipseq v2.1.0 ^88^ of the nf-core collection of workflows ^89^ with parameters ’--macs_gsize 2652783500 --narrow_peak --macs_fdr 0.01’. In brief, adapters and low-quality reads were removed using Trim Galore (v0.6.7), trimmed reads were mapped to the mouse reference genome (GRCm38, GENCODE vM15, primary assembly) using BWA ^90^. Reads mapping to blacklisted regions were removed, as well as those marked as duplicates (Picard v3.2.0), not uniquely mapped (SAMtools ’-F 0×004 -F 0×400 -q 1’), or containing > 4 mismatches. Narrow peaks were called by MACS3 (v 3.0.1) ^91^ for each sample separately (each with own input as control) and subsequently merged to form the final peakset. These peaks were annotated relative to gene features using HOMER’s annotatePeaks.pl (v 4.11) ^92^, and reads within them counted with featureCounts (v4.0.1) ^93^.

For the previously published ChIP-seq dataset from ^44^, FASTQ files were downloaded from GEO GSE116990 using nf-core/fetchngs, and reprocessed with nf-core/chipseq pipeline v2.1.0 with parameters (--macs_gsize 2652783500 --narrow_peak --macs_fdr 0.0001 -- min_reps_consensus 2). For H3K27ac and H3K4me1 this used the consensus peaks of 2 replicates against input (WT_0h), and for HDAC3 peaks this just used ChIPseq_FLAG_TX1072_Hdac3-AID-FLAG_0h_Dox_Rep1 against ChIPseq_FLAG_TX1072_WT_0h_Dox_Rep1. Consensus peaks for H3K27ac and H3K4me1 were further overlapped using BEDTools intersect ^94^, identifying 47,507 putative active regulatory elements. These were further classified based on presence of HDAC3 peaks within 100 bp distance (using bedtools window -w 100 -u) and motif analysis was performed using HOMER’s findMotifsGenome.pl (mm10 -size given -mask) on those with overlapping or nearby HDAC3 peaks versus those without.

### Transcriptomics and nascent transcriptomics

#### TT_chem_-seq

Analysis of nascent transcriptome was performed as described by Gregersen et al.^53^. Briefly, cells were labelled for 15 minutes with 1 mM of the nucleotide analogue 4-thiouridine (4sU), followed by RNA extraction using TRIzol and isopropanol precipitation. Extracted RNA (100 µg) was mixed to 1 µg of *Saccharomyces cerevisiae* RNA, labelled for 5 minutes with 5 mM 4-thiouracil (4tU). RNA mixture was fragmented with 0.2 M NaOH on ice for 20Lmin, neutralized, and purified using phenol:chloroform:isoamylalkohol extraction and isopropanol precipitation. Fragmented RNA was biotinylated using MTSEA-biotin-XX (Insight Biotechnologies, cat.no.: 90066, dissolved in DMF) and purified by phenol:chloroform extraction and isopropanol precipitation. Biotin-labeled RNA was enriched using streptavidin beads (Miltenyi μMACS), washed, and eluted with DTT. RNA was purified with RNeasy MinElute Cleanup kit (QIAGEN)columns and quality checked with TapeStation (Agilent). Libraries were prepared using the NEBNext Ultra II Directional RNA Kit and dual-index UMI adapters (NEB), following the rRNA-depleted FFPE RNA protocol. Single-end sequencing was performed in a P3-50bp kit on a NestSeq2000 sequencer.

#### Data processing

Raw sequencing data were demultiplexed and converted into FASTQ files using Illumina bcl2fastq Conversion Software (v2.20.0.422). UMI-tools extract (v 1.1.4) ^95^ was used to extract the UMI barcode from read2 and add it to the read name of read1, adapter sequences and low-quality bases were trimmed using Trim Galore. Reads were first mapped to a single copy of the ribosomal DNA repeat unit (GenBank: BK000964.3) to remove related transcribed sequences. Reads that did not map to rDNA were then mapped to yeast genome (sacCer3/R64 assembly), and the unmapped reads subsequently mapped to the mouse genome (GRCm38, GENCODE vM15, primary assembly) using STAR (v2.7.2d) ^96^. Resulting bam files were deduplicated using UMI-tools dedup, and lastly count matrices for genes (as annotated in GENCODE vM15) were generated using featureCounts (-s 2 -t gene).

To identify non-genic transcripts from nascent RNA sequencing reads, HOMER (v4.11) ^92^ was used to first combine deduplicated bam files from 2iLIF conditions (+/-IAA) into one tag directory and then run findPeaks on it with parameters ’-style groseq -rev -minBodySize 100’. Subsequently, all identified transcripts overlapping the sense strand of annotated genes (GENCODE vM15) or 5kb downstream were removed using bedtools window (v2.31.0) ^94^, and remaining transcripts within 100bp were merged (bedtools merge -s -d 100), resulting in 30,195 transcripts. Overlapping reads were counted with featureCounts (-s 2).

#### RNA-sequencing

RNA was extracted using RNAeasy Plus Mini kit with DNA digestion (QIAGEN). RNA integrity was assessed using Fragment Analyzer (AATI). For library preparation we used the Universal Plus™ Total RNA-Seq with NuQuant^®^ (TECAN) following manufacturer’s instructions. Single-end sequencing was performed using Illumina P3-50bp kit on a NextSeq2000 sequencer.

### Mouse breeding, embryo collection and dissection

Timed natural matings were used for all experiments. Noon of the day when the vaginal plugs of mated females were identified was scored as E0.5. A published conditional allele of *Hdac3^loxP^* was used ^97^. To obtain *Hdac3^Het^* or *Hdac3^KO^* embryos, *Hdac3^loxP/loxP^ Zp3-Cre^+ve^* females were crossed with *Hdac3^+/-^* males ^74^. All work was carried out in accordance with European legislation, authorized by the Danish National Animal Experiments Inspectorate (Dyreforsøgstilsynet, license no. 2020-15-0201-00609 and 2020-15-0201-00608) and performed according to national guidelines.

### Flash-seq

RNA extraction was performed using the Arcturus PicoPure^®^ (Applied Biosystems) from E4.5 blastocysts or E6.5 dissected epiblasts according to Chenoweth and Tesar ^98^. RNA was eluted in 10 µL, 4 µL of which were used for library construction. Libraries were prepared as described here https://www.protocols.io/view/flash-seq-protocol-kxygxzkrwv8j/v3?version_warning=no ^79^ after volume adjustment. Paired-end sequencing was performed using P1-100bp & P2-100bp Illumina sequencing kit on a NextSeq2000 sequencer.

#### Embryo genotyping

Whole embryos were lysed in 10 µL of Genotyping Buffer (200 mM NaCl, 100 mM Tris-HCl, 0.5% Tween 20) + 0.8 mg/mL Proteinase K by incubation for 4 hours at 56°C followed by 20 minutes at 95°C. Equal volume of PCR-grade water was added to each sample and the lysates were genotyped by PCR, with primers amplifying the *Hdac3* locus, the *Zp3-cre* locus, and sex chromosome-linked loci. Alternatively, cDNA generated following the Flash-seq protocol was used as a template for Real Time RT-qPCR, with primers detecting the *Hdac3* wild-type or mutated allele. Sex distinction was performed by primers detecting the *Xist* RNA or the transcript of the Y-linked *Eif2s3y* gene.

#### Data processing

Raw sequencing data were demultiplexed and converted into FASTQ files using Illumina bcl2fastq Conversion Software (v2.20.0.422). FASTQ files were processed according to the FLASH-seq UMI protocol V.3 [https://www.protocols.io/view/flash-seq-umi-protocol-bp2l619rdvqe/v3?step=12], focusing on the internal reads only, with read counting done on the level of ENSEMBL gene IDs as opposed to gene symbols. In brief, UMI-containing reads were identified using umi_tools extract and filtered out using BBTools’ filterbyname.sh, trimmed using BBDuk, mapped using STAR (v2.7.2d) against GRCm38, with parameters as given in the protocol. Lastly a count matrix was generated using subread’s featureCount (v2.0.6) with annotations from GENCODE release M15 primary assembly (--fracOverlap 0.25).

### Data Analysis - Bioinformatics

We performed Differential Gene Expression Analysis using DESeq2 (1.46.0) ^99^. The variance-stabilized transformed counts that were used for the Principal Component Analysis (PCA) plot were corrected using limma R package (3.62.2) ^100^ for batch effects introduced for either the sequencing run (FLASH-seq) or the collection day (replicate - TT_chem_-seq and RNA-seq runs). Raw uncorrected counts were used as input for DESeq2, and the batch parameter was taken into account in the model. Moreover, we used independent filtering from the in-built DESeq2 function with alpha set at 0.05. Gene expression changes were expressed as shrunk log2(Fold changes) using lfcShrink (type=”normal”). Transcript levels were calculated as sizeFactor-normalized fragments per kilobase per million reads (FPKM) using the argument “robust=TRUE”. For Gene Set Enrichment analysis, we used the collapsePathways function (fgsea R package 1.32.0) ^101^ and the lists were ranked based on the stat value generated from DESeq2. Differential Expression (DE) analysis for HOMER-identified transcripts was performed with DESeq2 using size factors calculated from genic reads. DE analysis for ChIP-seq and for Proteomics was performed using the edgeR package in r (4.4.2) ^102^. Heatmaps were visualized using Easeq – Interactive Software for analysis and visualization of ChIP-seq data (https://easeq.net) ^103^.

## Extended Data Figure Legends

**Extended Data Fig 1. HDAC3 resides at highly active enhancers.**

**a–b**, Heatmaps of all putative regulatory elements (±5,000 bp) bound (**a**) or not bound (**b**) by HDAC3. Shown is enrichment of HDAC3-FLAG and no FLAG control as well as H3K4me1, H3K4me3, and H3K27ac. Data was downloaded from Ż*ylicz et al.* ^44^ and originates from female ESCs. Regions are sorted in descending order of HDAC3 signal. Color intensity indicates reads per million per 1 kb, binned at 100 bp intervals.

**c**, Bar plot showing genomic location of putative regulatory elements bound or not bound by HDAC3. Bar lengths represent percentages. Statistical significance was assessed using the chi-square test.

**d**, Violin plot showing H3K27ac ChIP signal normalized to input at putative regulatory elements bound or not bound by HDAC3. Ratios were calculated from RPKMs. Statistical analysis was performed using the Wilcoxon–Mann–Whitney test.

**e**, Proteins identified in P300 proximity ligation experiment in undifferentiated ESC. Shown are reanalyzed normalized LFQ intensities extracted from *Stelloo et al.* ^52^. Statistical analysis was performed using DESeq2.

**Extended Data Fig 2. HDAC3 regulates highly active enhancers and their nearby genes**.

**a**, Immunoblot analysis of whole-cell extracts from TIR1 only or HDAC3-mAID-FLAG degron ESCs treated with indole acetic acid (IAA) for varying durations. Blots were probed with anti-HDAC3, anti-FLAG, anti-LAMIN B1 and anti-VINCULIN antibodies.

**b**, Euler plot illustrating overlap of H3K27ac and H3K4me1 enriched putative regulatory elements with HOMER-transcript loci and/or HDAC3 peaks.

**c**, GSEA for HDAC3-bound reRNAs against a ranked list of all identified HOMER transcripts. Ranking was performed using the stat value from DESeq2.

**d**, GSEA of genes differentially expressed in ESCs in 2iLIF with or without IAA, for genes associated with GO terms of biological processes.

**e**, Violin plot showing the distance of differentially expressed genes (as in d) from the nearest HDAC3 peak, categorized as upregulated, downregulated, or unchanged. Statistical analysis was performed using Kruskal–Wallis rank sum test followed by Dunn’s test. *P*-values were corrected using the Benjamini–Hochberg method.

**f**, Quantification of transcriptional activity in ESCs treated with or without IAA for 3 hours, measured by EU incorporation and normalized to DAPI intensity. Statistical analysis was performed using the Mann–Whitney test on two experimental replicates and two independent clones (N=4).

**Extended Data Fig 3. HDAC3 catalytic activity is involved in its function *in cis*.**

**a**, Box plots showing transcription levels of genes upregulated upon HDAC3 loss in ESCs and quantified in ESCs treated with or without IAA, and complemented by HDAC3-WT, HDAC3-Y298F, or empty vector.

**b**, MA-plot showing the gene shrunk log2(fold change) between HDAC3-mAID-FLAG degron 2iLIF ESCs complemented with HDAC3^Y298F^ vs empty vector. X axis shows nascent transcription levels in unperturbed 2iLIF degron ESCs.

**c**, Violin plot showing nascent transcription levels in the untreated 2iLIF ESCs. Plotted are upregulated, downregulated and unchanged genes upon HDAC3^Y298F^ expression versus empty vector control. Transcript levels are size-factor normalized fragments per kilobase per million reads (FPKMs). Statistical analysis was performed using the Kruskal–Wallis rank sum test followed by Dunn’s test. *P*-values were corrected using the Benjamini–Hochberg method.

**d-e**, GO-term analysis of genes upregulated (d) or downregulated (e) as described in (**b**).

**f,** MA-plot as described in (**b**) illustrating differences between HDAC3-mAID-FLAG degron 2iLIF ESCs complemented with HDAC3^WT^ vs empty vector.

**g**, Violin plot showing nascent transcription levels in the untreated 2iLIF ESCs. Plotted are upregulated, downregulated and unchanged genes upon HDAC3^WT^ expression versus empty vector control. Transcript levels are size-factor normalized fragments per kilobase per million reads (FPKMs). Statistical analysis was performed using the Kruskal–Wallis rank sum test followed by Dunn’s test. *P*-values were corrected using the Benjamini–Hochberg method.

**h**, Euler plot showing the overlaps between genes upregulated or downregulated in the following conditions: HDAC3-mAID-FLAG degron 2iLIF ESCs ±IAA, HDAC3-mAID-FLAG degron 2iLIF ESCs complemented with HDAC3^WT^vs an empty vector or complemented with HDAC3^Y298F^ vs an empty vector.

**Extended Data Fig 4. HDAC3 role in enhancer regulation is specific to pluripotent cells.**

**a**, Immunofluorescence images of TSCs, stained for TSC markers CDX2 and EOMES, extraembryonic mesoderm marker T (BRACHYURY), and pluripotency marker POU5F1.

**b**, MA-plot of shrunk log2(fold change) for differentially expressed genes in IAA-treated versus untreated TSCs against nascent transcription in unperturbed degron TSCs.

**c**, GSEA of genes differentially expressed in TSCs in FAXY with or without IAA, for genes associated with GO terms of biological processes.

**d**, Venn diagram showing overlap of DE genes in TSCs±IAA and DE genes in ESCs treated with or without IAA for 3 hours.

**e**, Venn diagram showing overlap between the top quartile of most expressed genes in ESCs and TSCs.

**f**, Histogram of common highly expressed genes from (**e**) showing the distribution of log_2_(fold changes) after IAA treatment in ESCs and TSCs.

**Extended Data Fig 5. HDAC3 regulates highly active enhancers and their nearby genes.**

**a**, Volcano plot showing distribution of expression changes for DE genes in ESCs switched to FA medium for 3 hours.

**b**, Heatmap illustrating expression changes of selected genes as in (**a**).

**c**, Violin plot showing basal transcription levels of DE genes in ESCs induced with FA medium for 3 hours, with or without IAA. Statistical analysis was performed using the Kruskal–Wallis rank sum test followed by Dunn’s test. *P*-values were corrected using the Benjamini–Hochberg method.

**d**, GSEA of genes differentially expressed in FA-induced ESCs with or without IAA, for genes associated with GO terms of biological processes.

**Extended Data Fig 6. HDAC3 prevents exit from naïve pluripotency and capacitation.**

**a**, Heatmap illustrating expression changes of selected naïve, core and primed pluripotency markers in ESCs and EpiLCs.

**b**, Venn diagram showing the overlap of DE genes between identified by RNA-seq in the 48-hour EpiLC±IAA treatment and the DE genes identified by TT_chem_-seq in the 3-hour FA±IAA treatment.

**c-d**, Immunofluorescence images (**c**) and quantification (**d**) of ESCs induced into EpiLCs with or without IAA, stained for a naïve pluripotency marker TBX3 and core pluripotency marker POU5F1. Statistics were performed using the the Wilcoxon–Mann–Whitney test.

**e**, Schematic of the workflow for short reversion experiments after introduction of vectors encoding HDAC3-WT, HDAC3-Y298F, or the empty vector. Cells were induced in FA for 24 hours and reverted to 2iLIF for 24 hours.

**f**, Violin plot of ratios between bootstrapped medians of TBX3 immunofluorescence signal for ESCs that underwent short reversion. Bootstrapping was performed 5,000 times. Statistics used Kruskal–Wallis rank sum test followed by Dunn’s test. *P*-values were corrected using the Benjamini–Hochberg method.

**g**, Violin plots of per plate colony count in long term reversion experiments (as in Fig. 5h). Each dot represents an independent experimental replicate. Representative images of plates used for quantifications. Statistical analysis was performed using Kruskal–Wallis rank sum test followed by Dunn’s test. *P*-values were corrected using the Benjamini–Hochberg method.

**Extended Data Fig 7. HDAC3 prevents exit from naive pluripotency.**

**a**, Schematic of workflow for short reversion experiment: cells induced in FA for 24 hours, then reverted to 2iLIF for 24 hours.

**b–c**, Immunofluorescence images (c) of cells as in (a), and quantification of TBX3 signal normalized to DAPI intensity (b). Two experimental replicates are shown (top and bottom). Statistical analysis was performed using Mann-Whitney test.

**Extended Data Fig 8. HDAC3 is required for early mouse development.**

**a**, Representative wholemount immunofluorescence images of *Hdac3*-Het or *Hdac3*-KO mouse E6.5 embryos. DAPI signal is cyan; NANOG is magenta. Epiblast volume is highlighted in white. **b**, Quantification of the ratio of NANOG signal to epiblast volume, using Imaris Software. Statistical analysis was performed using unpaired t-test.

**c**, Volcano plot analysis of gene expression levels in E6.5 female *Hdac3-*KO vs. *Hdac3-*Het epiblasts with shrunk log_2_(fold change) and -log10(adjusted p-value) calculated using DESeq2. Differentially expressed genes with adjusted p-value<0.05 are highlighted in color.

**d**, Violin plot showing gene expression levels in the *Hdac3-*Het female E6.5 epiblasts. Plotted are upregulated, downregulated and unchanged genes in *Hdac3-*KO vs. *Hdac3-*Het E6.5 female epiblasts. Transcript levels are size-factor normalized fragments per kilobase per million reads (FPKMs). Statistical analysis was performed using the Kruskal–Wallis rank sum test followed by Dunn’s test. *P*-values were corrected using the Benjamini–Hochberg method.

**e**, GSEA of differentially expressed genes in E6.5 *Hdac3-*KO vs. *Hdac3-*Het female epiblasts for genes associated with GO terms of biological processes.

**f–g**, Venn diagrams showing overlap of DE genes changing in E6.5 *Hdac3-*KO vs. *Hdac3-*Het male E6.5 epiblasts and DE genes changing in 2iLIF±IAA and FA±IAA conditions, respectively. **h-i**, Volcano plot analysis of gene expression levels in E4.5 male (h) and female (i) *Hdac3-*KO vs. *Hdac3-*Het embryos with shrunk log_2_(fold change) and -log10(adjusted p-value) calculated using DESeq2. Differentially expressed genes with adjusted p-value<0.05 are highlighted in color. Shown are also violin plots quantifying gene expression levels in the *Hdac3-*Het male(h) and female(i) E4.5 embryos. Plotted are upregulated, downregulated and unchanged genes in *Hdac3-*KO vs. *Hdac3-*Het embryos. Transcript levels are size-factor normalized fragments per kilobase per million reads (FPKMs). Statistical analysis was performed using the Kruskal–Wallis rank sum test followed by Dunn’s test. *P*-values were corrected using the Benjamini–Hochberg method.

## Acknowledgments

This work was supported by grants from: Novo Nordisk Fonden [NNF21CC0073729(JJZ), NNF20OC0059959 (LHG)], Danmarks Frie Forskningsfond [0169-00031B(JJZ), 0134-00190B (JJZ), 0165-00092B(LHG)], Danish National Research Foundation [DNRF154(LHG)] and European Research Council [ChroMeta: 101077271(JJZ), TranscriptStress: 101076758(LHG)]. We are thankful to Dr François Dossin for reading and reviewing the manuscript, and also for sharing the HDAC3-mAID-FLAG vector. We are grateful to Prof. Mitchell Lazar for sharing the *Hdac3* mouse line. We are also specially grateful for the critical feedback of Joshua Brickman. We thank reNEW platforms for technical expertise, support, and use of equipment in particular: H. Wollmann, M. Michaut, J. Bulkescher, and A. Kalvisa. We also thank EMBL Rome core facilities, particularly Gerald Pfister (FCF). Language in some paragraphs was proof-read with Copilot and Perplexity; however no passage was written by and no references were added using large language models.

## Authors Contributions Statement

Conceptualization: N.S., J.J.Z.; Data curation: N.S., A.W., J.J.Z.; Formal analysis: N.S., A.W., J.J.Z.; Funding acquisition: L.H.G., J.A.H., J.J.Z.; Investigation (omics experiments): N.S., S.K., G.N. K.K.U., A.W. J.J.Z..; Investigation (*in vivo* experiments): N.S., S.B.A., J.J.Z.; Investigation (stem cell experiments): N.S.; Project administration: J.J.Z; Resources: L.H.G., J.J.Z.; Software: N.S., K.K.U., A.W.; Supervision: L.H.G., J.A.H., J.J.Z; Validation: N.S.; Visualization: N.S.; Writing, reviewing and editing of the manuscript: N.S., A.W, K.K.U., L.H.G., J.A.H., J.J.Z.

