## Supplementary figures and images for "HDAC3 prevents enhancer hyperactivation to enable developmental transitions"

### Extended Data Figure 1

#### Extended Data Figure 1

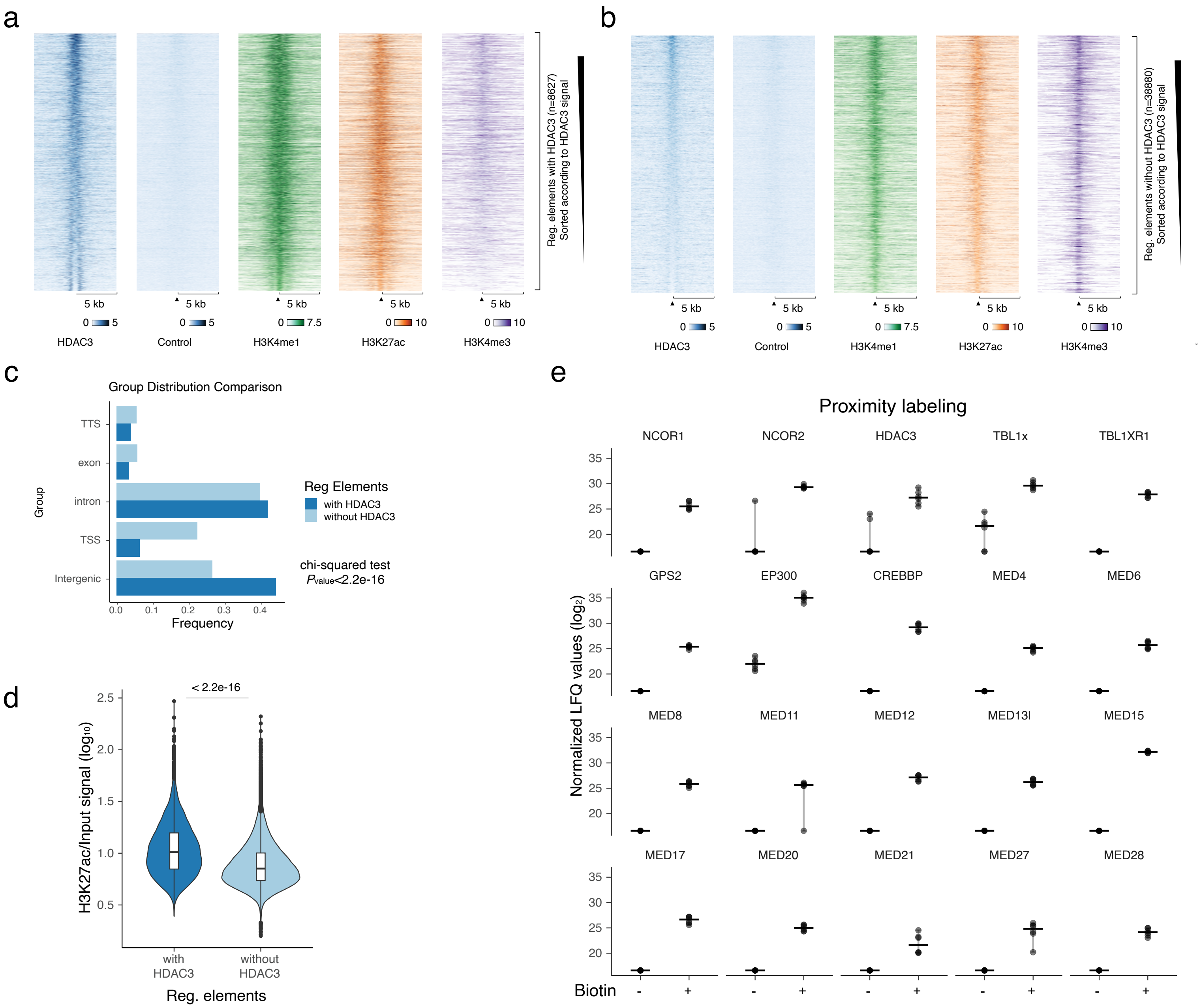

### Extended Data Figure 2

#### Extended Data Figure 2

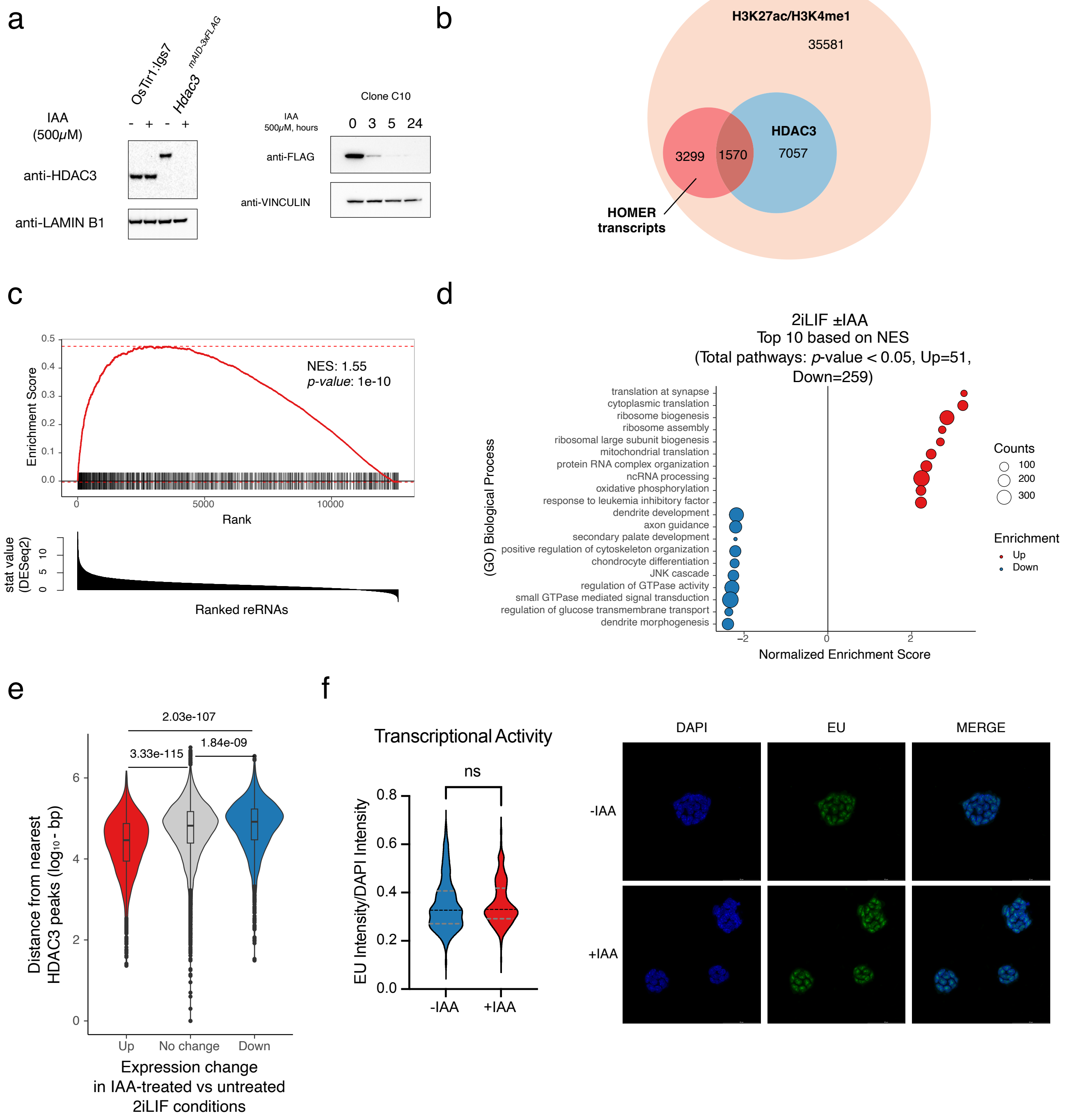

### Extended Data Figure 3

#### Extended Data Figure 3

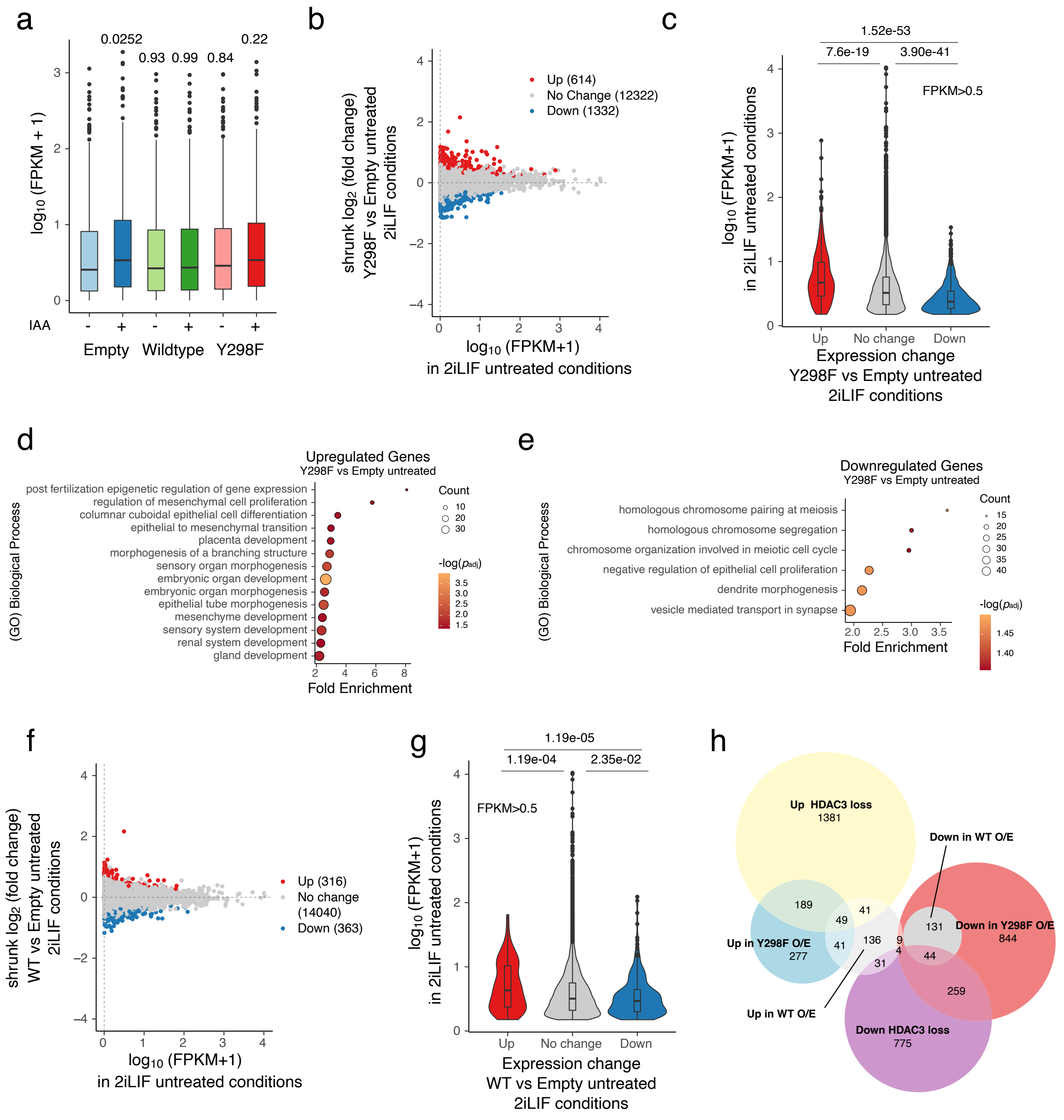

### Extended Data Figure 4

#### Extended Data Figure 4

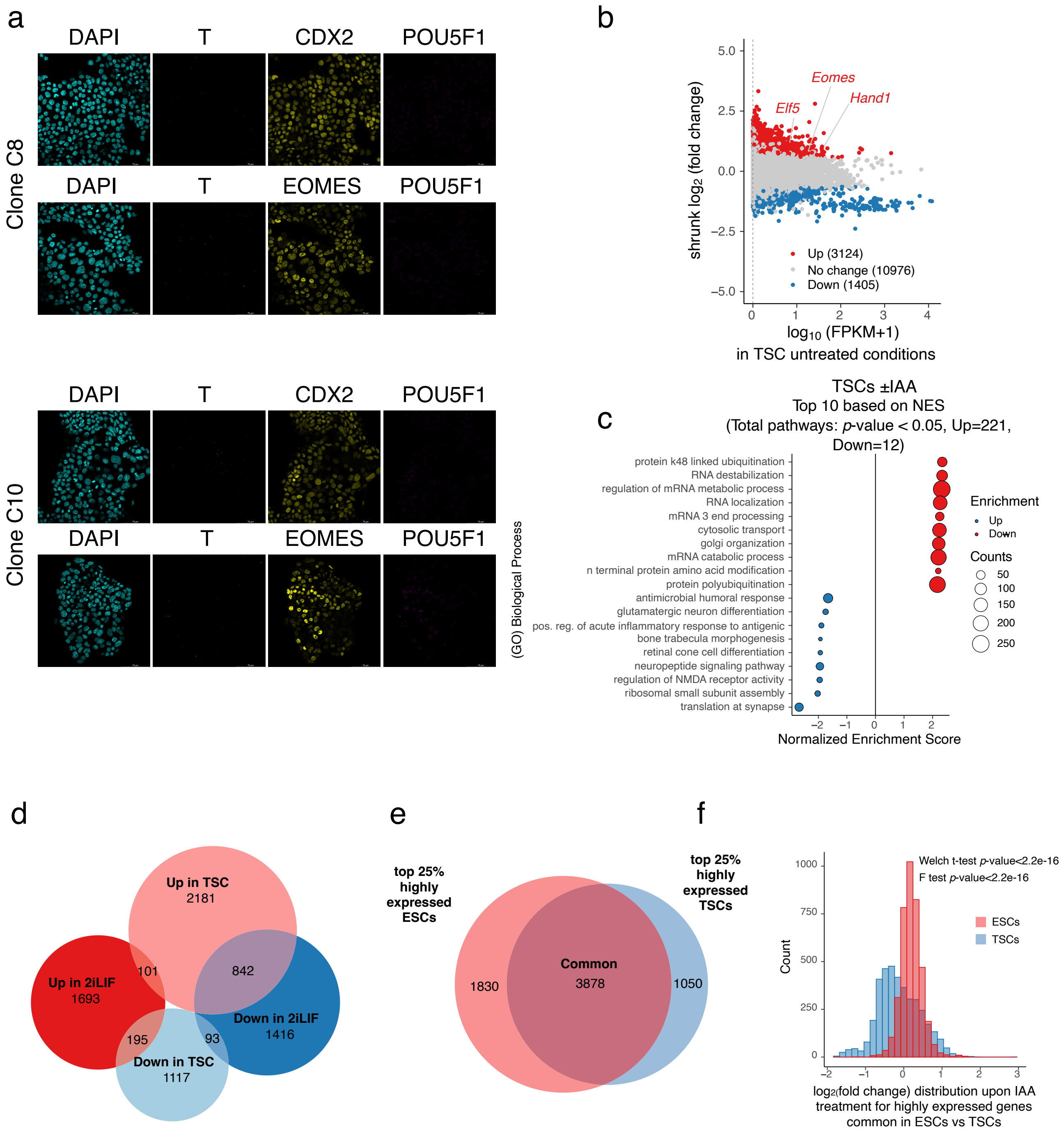

### Extended Data Figure 5

#### Extended Data Figure 5

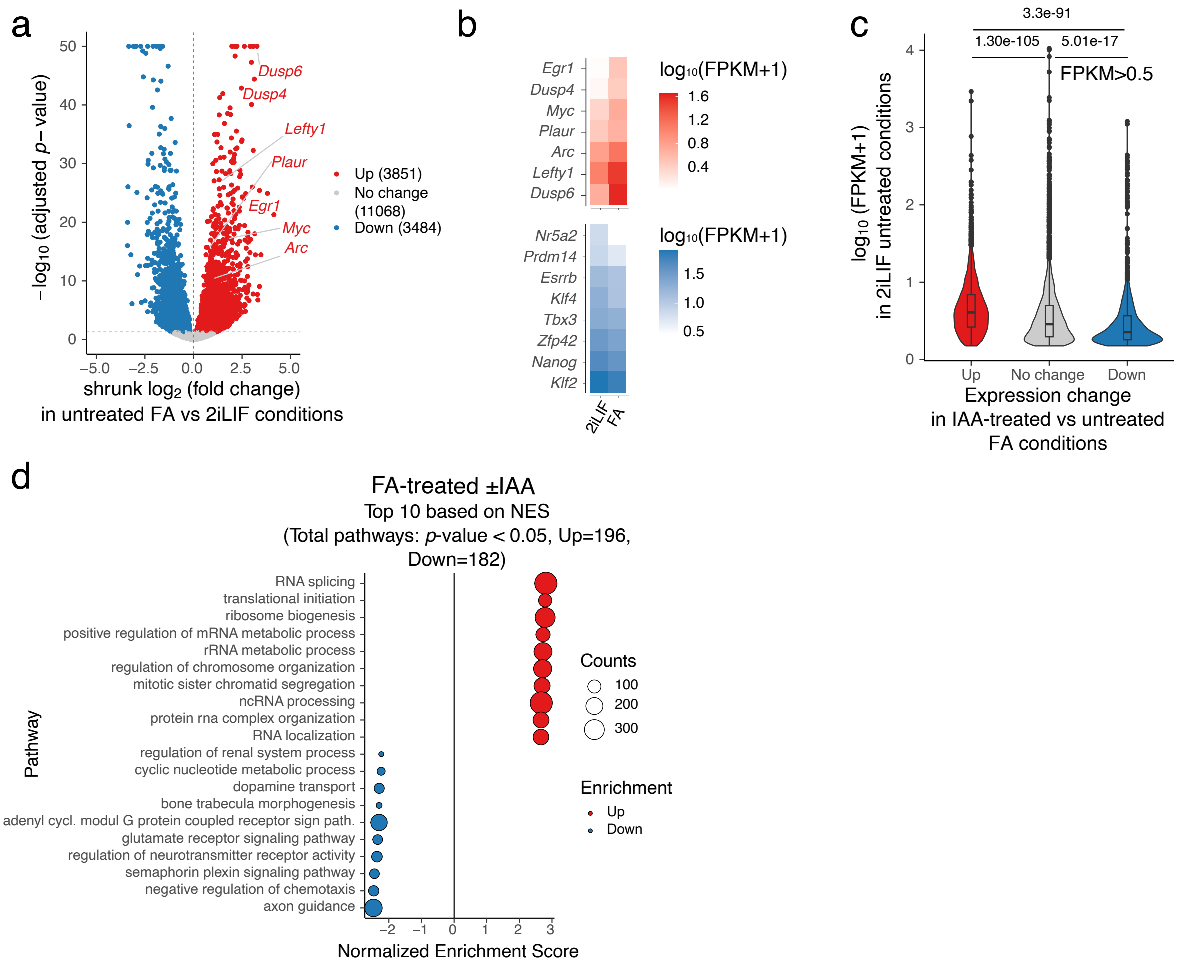

### Extended Data Figure 6

#### Extended Data Figure 6

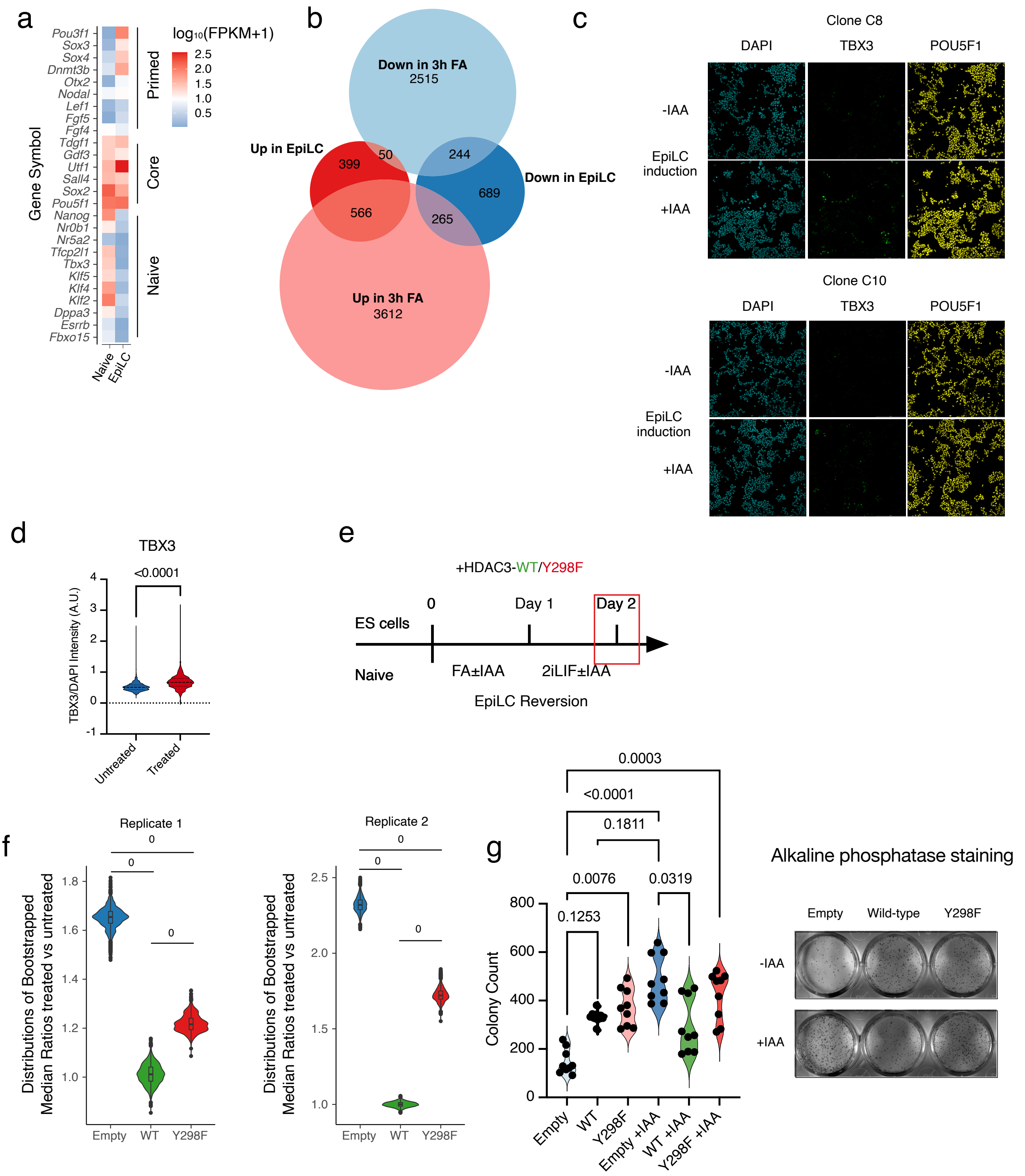

### Extended Data Figure 7

#### Extended Data Figure 7

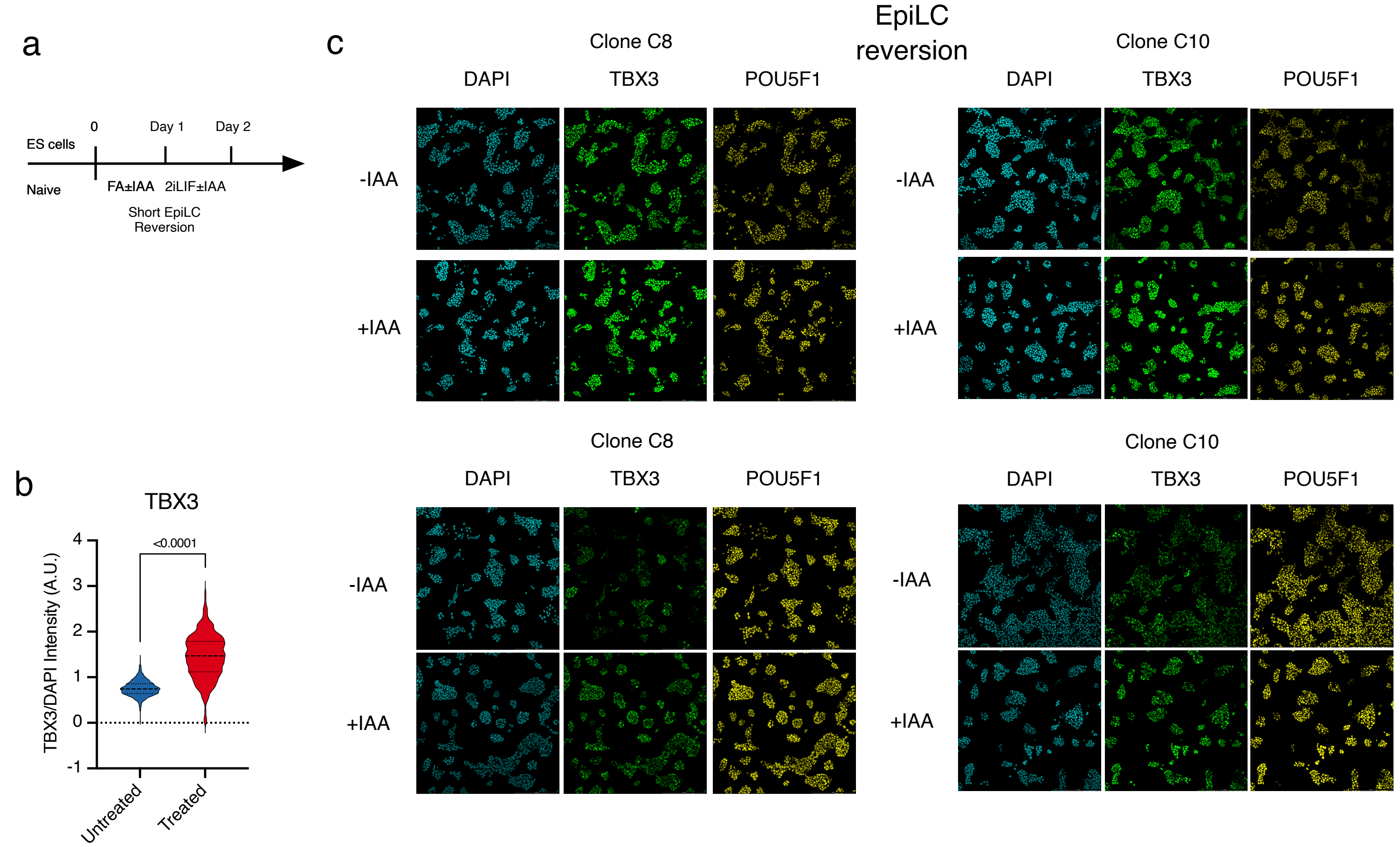

### Extended Data Figure 8

#### Extended Data Figure 8

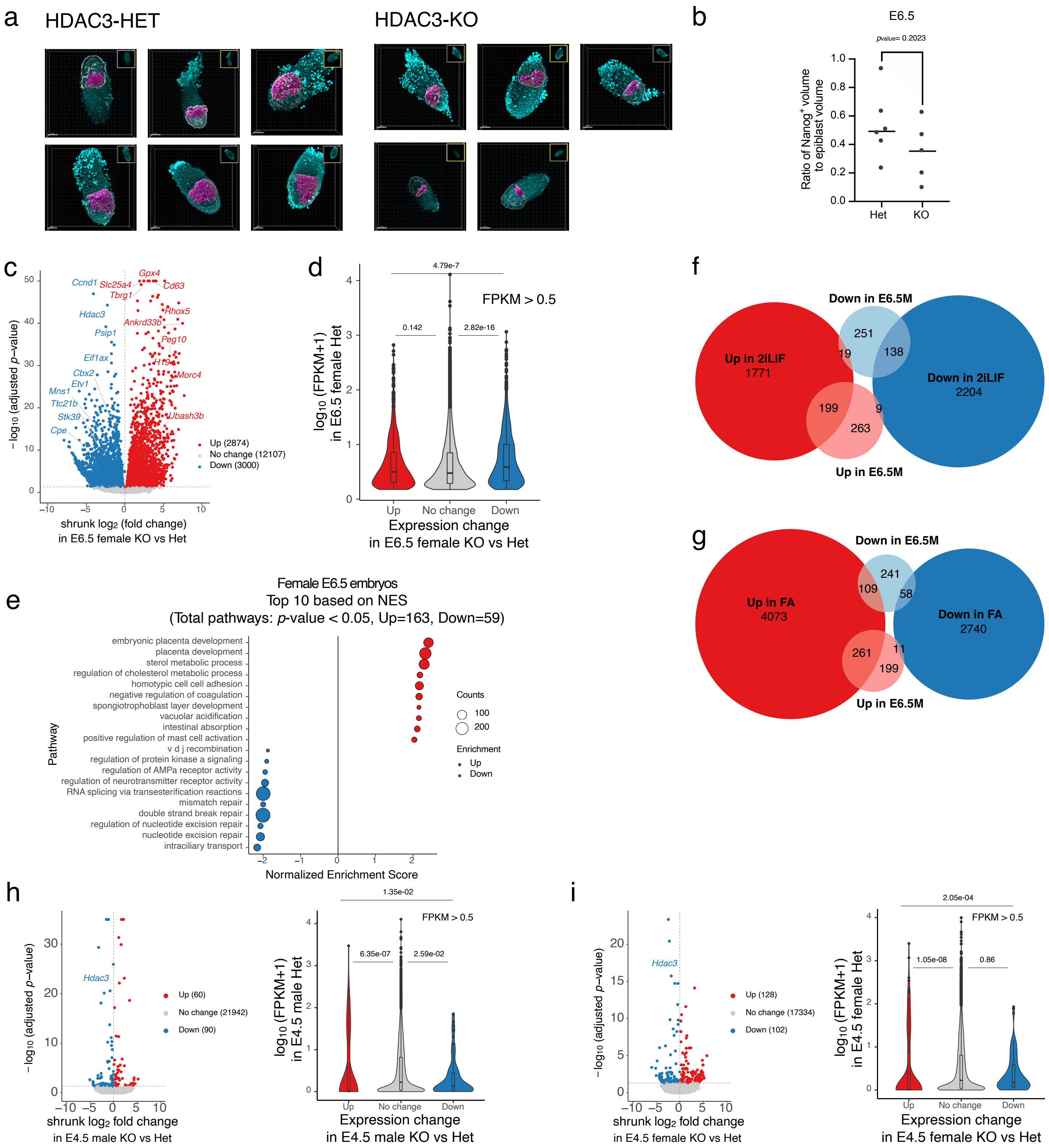
