## Supplementary Table 3 for "HDAC3 prevents enhancer hyperactivation to enable developmental transitions"

**FLASH-seq E4.5 Male**

out of 24931 with nonzero total read count
adjusted p-value < 0.05
LFC > 0 (up) : 60, 0.24%
LFC < 0 (down) : 90, 0.36%
outliers [1] : 269, 1.1%
low counts [2] : 0, 0%
(mean count < 0)
[1] see 'cooksCutoff' argument of ?results
[2] see 'independentFiltering' argument of ?results

**FLASH-seq E4.5 Female**

out of 24931 with nonzero total read count
adjusted p-value < 0.05
LFC > 0 (up) : 128, 0.51%
LFC < 0 (down) : 102, 0.41%
outliers [1] : 269, 1.1%
low counts [2] : 6617, 27%
(mean count < 2)
[1] see 'cooksCutoff' argument of ?results
[2] see 'independentFiltering' argument of ?results

**FLASH-seq E6.5 Male**

out of 24931 with nonzero total read count
adjusted p-value < 0.05
LFC > 0 (up) : 471, 1.9%
LFC < 0 (down) : 408, 1.6%
outliers [1] : 269, 1.1%
low counts [2] : 11350, 46%
(mean count < 12)
[1] see 'cooksCutoff' argument of ?results
[2] see 'independentFiltering' argument of ?results

Kruskal-Wallis rank sum test

data: fpkm by group
Kruskal-Wallis chi-squared = 489.59, df = 2, p-value < 2.2e-16

Dunn Test

| .y. | group1 | group2 | n1 | n2 | statistic | p | p.adj |
|---|---|---|---|---|---|---|---|
| fpkm | up | noDE | 448 | 11862 | -21.94 | 9.92e-107 | 2.98e-106 |
| fpkm | up | down | 448 | 374 | -12.40 | 2.53e-35 | 3.79e-35 |
| fpkm | noDE | down | 11862 | 374 | 3.57 | 3.58e-04 | 3.58e-04 |

**FLASH-seq E6.5 Female**

out of 24931 with nonzero total read count
adjusted p-value < 0.05
LFC > 0 (up) : 2874, 12%
LFC < 0 (down) : 3000, 12%
outliers [1] : 269, 1.1%
low counts [2] : 6617, 27%
(mean count < 2)
[1] see 'cooksCutoff' argument of ?results
[2] see 'independentFiltering' argument of ?results

Kruskal-Wallis rank sum test

data: fpkm by group
Kruskal-Wallis chi-squared = 171.32, df = 2, p-value < 2.2e-16

| .y. | group1 | group2 | n1 | n2 | statistic | p | p.adj |
|---|---|---|---|---|---|---|---|
| fpkm | up | noDE | 2098 | 7931 | -6.60 | 4.02e-11 | 6.03e-11 |
| fpkm | up | down | 2098 | 2607 | 4.10 | 4.14e-05 | 4.14e-05 |
| fpkm | noDE | down | 7931 | 2607 | 12.5 | 6.84e-36 | 2.05e-35 |

**TTchem-seq FA untreated vs 2iLIF untreated**

out of 22838 with nonzero total read count
adjusted p-value < 0.05
LFC > 0 (up) : 4719, 21%
LFC < 0 (down) : 5722, 25%
outliers [1] : 0, 0%
low counts [2] : 1329, 5.8%
(mean count < 1)
[1] see 'cooksCutoff' argument of ?results
[2] see 'independentFiltering' argument of ?results

**TTchem-seq 2iLIF treated vs 2iLIF untreated**

out of 22838 with nonzero total read count
adjusted p-value < 0.05
LFC > 0 (up) : 1989, 8.7%
LFC < 0 (down) : 2351, 10%
outliers [1] : 0, 0%
low counts [2] : 6199, 27%
(mean count < 6)
[1] see 'cooksCutoff' argument of ?results
[2] see 'independentFiltering' argument of ?results

Kruskal-Wallis rank sum test

data:  fpkm by class
Kruskal-Wallis chi-squared = 2204.7, df = 2, p-value < 2.2e-16

| .y. | group1 | group2 | n1 | n2 | statistic | p | p.adj |
|---|---|---|---|---|---|---|---|
| fpkm | up | noDE | 1681 | 8412 | -31.14 | 7.52e-213 | 1.13e-212 |
| fpkm | up | down | 1681 | 1165 | -42.60 | 0.0e+00 | 0.0e+00 |
| fpkm | noDE | down | 8412 | 1165 | -25.34 | 1.16e-141 | 1.16e-141 |

**TTchem-seq FA treated vs FA untreated**

out of 22838 with nonzero total read count
adjusted p-value < 0.05
LFC > 0 (up) : 4443, 19%
LFC < 0 (down) : 2809, 12%
outliers [1] : 0, 0%
low counts [2] : 3100, 14%
(mean count < 2)
[1] see 'cooksCutoff' argument of ?results
[2] see 'independentFiltering' argument of ?results

Kruskal-Wallis rank sum test

data:  fpkm by class
Kruskal-Wallis chi-squared = 4895.8, df = 2, p-value < 2.2e-16

| .y. | group1 | group2 | n1 | n2 | statistic | p | p.adj |
|---|---|---|---|---|---|---|---|
| fpkm | up | noDE | 4171 | 6149 | -21.88 | 4.34e-106 | 1.30e-105 |
| fpkm | up | down | 4171 | 937 | -20.27 | 2.20e-91 | 3.3e-91 |
| fpkm | noDE | down | 6149 | 937 | -8.39 | 5.01e-17 | 5.01e-17 |

**Comparison of lists of DE genes: upregulated in 2iLIF±IAA vs upregulated in E6.5 Male Het vs KO**

Hypergeometric test: p-value = 1.351894e-87

Fisher's Exact Test for Count Data

data:  matrix_2iLIF_data_up
p-value < 2.2e-16
alternative hypothesis: true odds ratio is not equal to 1
95 percent confidence interval:
 6.922003 10.197469
sample estimates:
odds ratio
 8.408745

**Comparison of lists of DE genes: downregulated in 2iLIF±IAA vs downregulated in E6.5 Male Het vs KO**

Hypergeometric test: p-value = 1.364425e-38

Fisher's Exact Test for Count Data

data:  matrix_2iLIF_data_down
p-value < 2.2e-16
alternative hypothesis: true odds ratio is not equal to 1
95 percent confidence interval:
 3.757043 5.781977
sample estimates:
odds ratio
  4.668851

**Comparison of lists of DE genes: upregulated in FA±IAA vs upregulated in E6.5 Male Het vs KO**

Hypergeometric test: p-value =  1.439286e-68

Fisher's Exact Test for Count Data

data:  matrix_FA_data_up
p-value < 2.2e-16
alternative hypothesis: true odds ratio is not equal to 1
95 percent confidence interval:
 4.474323 6.533565
sample estimates:
odds ratio
  5.403905

**Comparison of lists of DE genes: downregulated in FA±IAA vs downregulated in E6.5 Male Het vs KO**

Hypergeometric test: p-value = 0.1336287

Fisher's Exact Test for Count Data

data:  matrix_FA_data_down
p-value = 0.2244
alternative hypothesis: true odds ratio is not equal to 1
95 percent confidence interval:
 0.8793208 1.5738003
sample estimates:
odds ratio
    1.1854

**GSEA: upregulated in FA±IAA against sorted induction list**

| ID | setSize | enrichmentScore | NES | pvalue | p.adjust | qvalue | rank |
|---|---|---|---|---|---|---|---|
| up | 4443 | -0.7051069 | -3.750867 | 1e-10 | 1e-10 | NA | 5848 |

**GSEA: downregulated in FA±IAA against sorted induction list**

| ID | setSize | enrichmentScore | NES | pvalue | p.adjust | qvalue | rank |
|---|---|---|---|---|---|---|---|
| Down | 2809 | 0.5559604 | 3.395896 | 1e-10 | 1e-10 | NA | 5499 |

**TTchem-seq Enhancers 2iLIF±IAA**

```
out of 30195 with nonzero total read count
adjusted p-value < 0.05
LFC > 0 (up)       : 2181, 7.2%
LFC < 0 (down)     : 9, 0.03%
outliers [1]       : 0, 0%
low counts [2]     : 17562, 58%
(mean count < 7)
[1] see 'cooksCutoff' argument of ?results
[2] see 'independentFiltering' argument of ?results
```

```
Asymptotic Two-Sample Brown-Mood Median Test

data:  fpkm by class (up, noDE)
Z = 9.9111, p-value < 2.2e-16
alternative hypothesis: true mu is not equal to 0

Asymptotic Wilcoxon-Mann-Whitney Test

data:  mean_fpkm by class (up, noDE)
Z = 9.8827, p-value < 2.2e-16
alternative hypothesis: true mu is not equal to 0
```

**Distance of genes from proximal DE enhancers**

For DE genes in 2iLIF±IAA

Kruskal-Wallis rank sum test

data:  distance_to_enhancer by class
Kruskal-Wallis chi-squared = 1143.4, df = 2, p-value < 2.2e-16

Dunn Test

| .y. | group1 | group2 | n1 | n2 | statistic | p | p.adj |
|---|---|---|---|---|---|---|---|
| dist_to_enhancer | up | noDE | 1989 | 18466 | 28.99 | 8.16e-185 | 1.22e-184 |
| dist_to_enhancer | up | down | 1989 | 2351 | 32.71 | 9.90e-235 | 2.97e-234 |
| dist_to_enhancer | noDE | down | 18466 | 2351 | 14.26776 | 3.48e-46 | 3.48-46 |

For DE genes in FA±IAA

Kruskal-Wallis rank sum test

data:  distance_to_enhancer by class
Kruskal-Wallis chi-squared = 309.77, df = 2, p-value < 2.2e-16

Dunn Test

| .y. | group1 | group2 | n1 | n2 | statistic | p | p.adj |
|---|---|---|---|---|---|---|---|
| dist_to_enhancer | up | noDE | 4443 | 15575 | 16.60 | 6.31e-62 | 1.89e-61 |
| dist_to_enhancer | up | down | 4443 | 2809 | 13.99 | 1.77e-44 | 2.660e-44 |
| dist_to_enhancer | noDE | down | 15575 | 2809 | 2.67 | 7.49e-03 | 7.49e-03 |

**ChIP-seq Comparison: HDAC3 signal at peaks which show differential binding in 2iLIF conditions.**

For C8 clone

| .y. | group1 | group2 | n1 | n2 | statistic | p | p.adj |
|---|---|---|---|---|---|---|---|
| rpkm | Down_2iLIF | Down_FAK | 875 | 875 | -31.30 | 5.26E-215 | 2.63E-214 |
| rpkm | Down_2iLIF | No_change 2iLIF | 875 | 8421 | -18.25 | 2.24E-74 | 3.05E-74 |
| rpkm | Down_2iLIF | No_change FAK | 875 | 8421 | -13.74 | 5.71E-43 | 7.13E-43 |
| rpkm | Down_2iLIF | Up_2iLIF | 875 | 1040 | -27.63 | 4.36E-168 | 1.09E-167 |
| rpkm | Down_2iLIF | Up_FAK | 875 | 1040 | 8.00 | 1.28E-15 | 1.37E-15 |
| rpkm | Down_FAK | No_change 2iLIF | 875 | 8421 | 23.88 | 5.03E-126 | 9.43E-126 |
| rpkm | Down_FAK | No_change FAK | 875 | 8421 | 28.38 | 3.26E-177 | 9.77E-177 |
| rpkm | Down_FAK | Up_2iLIF | 875 | 1040 | 4.98 | 6.27E-07 | 6.27E-07 |
| rpkm | Down_FAK | Up_FAK | 875 | 1040 | 40.61 | 0.00E+00 | 0.00E+00 |
| rpkm | No_change 2iLIF | No_change FAK | 8421 | 8421 | 10.38 | 3.04E-25 | 3.51E-25 |

For C10 clone

| .y. | group1 | group2 | n1 | n2 | statistic | p | p.adj |
|---|---|---|---|---|---|---|---|
| rpkm | Down_2iLIF | Down_FAK | 875 | 875 | -38.68 | 0 | 0 |
| rpkm | Down_2iLIF | No_change 2iLIF | 875 | 8421 | -18.74 | 2.2E-78 | 2.7E-78 |
| rpkm | Down_2iLIF | No_change FAK | 875 | 8421 | -22.81 | 3.9E-115 | 5.9E-115 |
| rpkm | Down_2iLIF | Up_2iLIF | 875 | 1040 | -33.30 | 4.0E-243 | 1.2E-242 |
| rpkm | Down_2iLIF | Up_FAK | 875 | 1040 | 5.36 | 8.6E-08 | 8.6E-08 |
| rpkm | Down_FAK | No_change 2iLIF | 875 | 8421 | 33.32 | 1.9E-243 | 7.1E-243 |
| rpkm | Down_FAK | No_change FAK | 875 | 8421 | 29.26 | 3.6E-188 | 7.7E-188 |
| rpkm | Down_FAK | Up_2iLIF | 875 | 1040 | 7.01 | 2.3E-12 | 2.5E-12 |
| rpkm | Down_FAK | Up_FAK | 875 | 1040 | 45.67 | 0.0E+00 | 0.0E+00 |
| rpkm | No_change 2iLIF | No_change FAK | 8421 | 8421 | -9.37 | 7.5E-21 | 8.7E-21 |

**GSEA: genes in proximity to Gained peaks against sorted FA induction genes.**

| ID | setSize | enrichmentScore | NES | pvalue | p.adjust | qvalue | rank |
|---|---|---|---|---|---|---|---|
| Gained | 546 | 0.2934345 | 1.643992 | 3.00e-08 | 3.00e-08 | NA | 4865 |

**GSEA: genes in proximity to Lost peaks against sorted FA induction genes.**

| ID | setSize | enrichmentScore | NES | pvalue | p.adjust | qvalue | rank |
|---|---|---|---|---|---|---|---|
| Lost | 517 | -0.38 | -1.86 | 1e-10 | 1e-10 | NA | 4120 |

**Distribution of HDAC3 peaks vs Regulatory Elements across the genome**

Chi-squared test for given probabilities

data:  HDAC3_total
X-squared = 1755.4, df = 5, p-value < 2.2e-16

[1] -11.304  27.073   3.476 -28.827  -7.165   0.468

Levels: ”Intergenic","promoter-TSS", "intron","exon","TTS", "NA"

**Distribution of Regulatory Elements with vs without HDAC3 across the genome**

Chi-squared test for given probabilities

data:  HDAC3_bound
X-squared = 2128.3, df = 5, p-value < 2.2e-16

[1]  -9.330  31.825   3.257 -31.318  -5.970   1.181

Levels: ”Intergenic","promoter-TSS", "intron","exon","TTS", "NA"

**Distance of DE genes in 2iLIF±IAA from HDAC3 peaks**

Kruskal-Wallis rank sum test

data:  distance_to_peak by class
Kruskal-Wallis chi-squared = 594.17, df = 2, p-value < 2.2e-16

Dunn Test

| .y. | group1 | group2 | n1 | n2 | statistic | p | p.adj |
|---|---|---|---|---|---|---|---|
| distance_to_peak | up | noDE | 1986 | 18428 | 22.86 | 1.11e-115 | 3.33e-115 |
| distance_to_peak | up | down | 1986 | 2350 | 22.03 | 1.36e-107 | 2.03e-107 |
| distance_to_peak | noDE | down | 18428 | 2350 | 6.01 | 1.84e-09 | 1.84-09 |

**Distance of DE genes in FA±IAA from HDAC3 peaks**

Kruskal-Wallis rank sum test

data:  distance_to_peak by class
Kruskal-Wallis chi-squared = 81.475, df = 2, p-value < 2.2e-16

Dunn Test

| .y. | group1 | group2 | n1 | n2 | statistic | p | p.adj |
|---|---|---|---|---|---|---|---|
| distance_to_peak | up | noDE | 4435 | 15545 | 8.46 | 2.78-17 | 8.34e-17 |
| distance_to_peak | up | down | 4435 | 2805 | 2.00 | 4.53e-02 | 4.53e-02 |
| distance_to_peak | noDE | down | 15545 | 2805 | -4.66 | 3.12e-06 | 4.68e-06 |

**TTchem-seq TSC treated vs TSC untreated**

out of 19708 with nonzero total read count
adjusted p-value < 0.05
LFC > 0 (up) : 3124, 16%
LFC < 0 (down) : 1405, 7.1%
outliers [1] : 0, 0%
low counts [2] : 4203, 21%
(mean count < 8)
[1] see 'cooksCutoff' argument of ?results
[2] see 'independentFiltering' argument of ?results

Kruskal-Wallis rank sum test

data:  fpkm by class
Kruskal-Wallis chi-squared = 662.75, df = 2, p-value < 2.2e-16

Dunn Test

| .y. | group1 | group2 | n1 | n2 | statistic | p | p.adj |
|---|---|---|---|---|---|---|---|
| fpkm | up | noDE | 3124 | 15179 | -5.06 | 4.22e-07 | 4.22e-07 |
| fpkm | up | down | 3124 | 1405 | 19.19 | 4.64e-82 | 6.96e-82 |
| fpkm | noDE | down | 15179 | 1405 | 25.67 | 2.71e-145 | 8.14=e-145 |

**Comparison of lists of DE genes in 2iLIF±IAA vs TSC±IAA**

Hypergeometric tests:

Up in 2iLIF±IAA vs Up in TSC±IAA: p-value = 1
Down in 2iLIF±IAA vs down in TSC±IAA: p-value = 0.9999969
Up in 2iLIF±IAA vs Down in TSC±IAA: p-value = 7.517203e-13
Down in 2iLIF±IAA vs Up in TSC±IAA: p-value = 5.600974e-197

Fisher's Exact Test for Count Data

data:  matrix_data_up (Up in 2iLIF±IAA vs Up in TSC±IAA)
p-value < 2.2e-16
alternative hypothesis: true odds ratio is not equal to 1
95 percent confidence interval:
 0.2675360 0.4065186
sample estimates:
odds ratio
 0.3313338

Fisher's Exact Test for Count Data

data:  matrix_data_down (Down in 2iLIF±IAA vs down in TSC±IAA)
p-value = 9.413e-06
alternative hypothesis: true odds ratio is not equal to 1
95 percent confidence interval:
 0.5027641 0.7818466
sample estimates:
odds ratio
 0.6301207

Fisher's Exact Test for Count Data

data:  matrix_data_mix1 (Up in 2iLIF±IAA vs Down in TSC±IAA)
p-value = 1.173e-12
alternative hypothesis: true odds ratio is not equal to 1
95 percent confidence interval:
 1.565640 2.164927
sample estimates:
odds ratio
 1.844809

Fisher's Exact Test for Count Data

data:  matrix_data_mix2 (Down in 2iLIF±IAA vs Up in TSC±IAA)
p-value < 2.2e-16
alternative hypothesis: true odds ratio is not equal to 1
95 percent confidence interval:
 4.242138 5.140317

sample estimates:
odds ratio
4.670334

**Comparison of common highly expressed genes in TSCs vs 2iLIF.**

F test to compare two variances

data: DE_TSC_genes$log2FoldChange[DE_TSC_genes$Row.names %in% common_high]
and DE_2iLIF_genes$log2FoldChange[DE_2iLIF_genes$Row.names %in% common_high]
F = 4.2406, num df = 3877, denom df = 3877, p-value < 2.2e-16
alternative hypothesis: true ratio of variances is not equal to 1
95 percent confidence interval:
3.981857 4.516226
sample estimates:
ratio of variances
4.240633

**TTchem-seq Empty treated vs untreated**

out of 23304 with nonzero total read count
adjusted p-value < 0.05
LFC > 0 (up) : 1715, 7.4%
LFC < 0 (down) : 1199, 5.1%
outliers [1] : 0, 0%
low counts [2] : 7681, 33%
(mean count < 8)
[1] see 'cooksCutoff' argument of ?results
[2] see 'independentFiltering' argument of ?results

**TTchem-seq WT treated vs untreated**

out of 23304 with nonzero total read count
adjusted p-value < 0.05
LFC > 0 (up) : 4, 0.017%
LFC < 0 (down) : 9, 0.039%
outliers [1] : 0, 0%
low counts [2] : 11295, 48%
(mean count < 36)
[1] see 'cooksCutoff' argument of ?results
[2] see 'independentFiltering' argument of ?results

**TTchem-seq Y298F treated vs untreated**

out of 23304 with nonzero total read count
adjusted p-value < 0.05
LFC > 0 (up) : 322, 1.4%
LFC < 0 (down) : 246, 1.1%
outliers [1] : 0, 0%
low counts [2] : 8585, 37%
(mean count < 12)
[1] see 'cooksCutoff' argument of ?results
[2] see 'independentFiltering' argument of ?results

**TTchem-seq Y298F untreated vs Empty untreated**

out of 23304 with nonzero total read count
adjusted p-value < 0.05
LFC > 0 (up) : 614, 2.6%
LFC < 0 (down) : 1332, 5.7%
outliers [1] : 0, 0%
low counts [2] : 9036, 39%
(mean count < 14)
[1] see 'cooksCutoff' argument of ?results
[2] see 'independentFiltering' argument of ?results

**TTchem-seq WT untreated vs Empty untreated**

out of 23304 with nonzero total read count
adjusted p-value < 0.05
LFC > 0 (up) : 316, 1.4%
LFC < 0 (down) : 363, 1.6%
outliers [1] : 0, 0%
low counts [2] : 8585, 37%
(mean count < 12)
[1] see 'cooksCutoff' argument of ?results
[2] see 'independentFiltering' argument of ?results

**TTchem-seq Log2FC across rescue cell lines (empty±IAA, WT±IAA, Y298F±IAA)**

Kruskal-Wallis rank sum test

data: log2FC by condition
Kruskal-Wallis chi-squared = 704.85, df = 2, p-value < 2.2e-16

Dunn Test

| .y. | group1 | group2 | n1 | n2 | statistic | p | p.adj |
|---|---|---|---|---|---|---|---|
| log2FC | empty | wildtype | 721 | 721 | -25.87 | 1.38e-147 | 4.15e-147 |
| log2FC | empty | Y298F | 721 | 721 | -7.78 | 7.47e-15 | 7.47e-15 |
| log2FC | wildtype | Y298F | 721 | 721 | 18.10 | 3.45e-73 | 5.18e-73 |

**TTchem-seq Y298F untreated vs Empty untreated**

Kruskal-Wallis rank sum test

data: fpkm by class
Kruskal-Wallis chi-squared = 274.8, df = 2, p-value < 2.2e-16

Dunn Test

| .y. | group1 | group2 | n1 | n2 | statistic | p | p.adj |
|---|---|---|---|---|---|---|---|
| fpkm | Up | noDE | 420 | 9963 | -8.87 | 7.60e-19 | 7.60e-19 |
| fpkm | Up | Down | 420 | 855 | -15.48 | 5.07e-54 | 1.52e-53 |
| fpkm | noDE | Down | 9963 | 855 | -13.48 | 1.95e-41 | 3.90e-41 |

**TTchem-seq WT untreated vs Empty untreated**

Kruskal-Wallis rank sum test

data:  fpkm by class
Kruskal-Wallis chi-squared = 21.591, df = 2, p-value = 2.049e-05

| .y. | group1 | group2 | n1 | n2 | statistic | p | p.adj |
|---|---|---|---|---|---|---|---|
| fpkm | Up | noDE | 150 | 10817 | -4.01 | 5.96e-05 | 1.19e-04 |
| fpkm | Up | Down | 150 | 271 | -4.61 | 3.98e-06 | 1.19e-05 |
| fpkm | noDE | Down | 10817 | 271 | -2.27 | 2.35e-02 | 2.34e-02 |

**RNA-seq EpiLC untreated vs Naive untreated**

out of 27456 with nonzero total read count
adjusted p-value < 0.05
LFC > 0 (up)       : 5357, 20%
LFC < 0 (down)     : 4922, 18%
outliers [1]       : 0, 0%
low counts [2]     : 5323, 19%
(mean count < 1)
[1] see 'cooksCutoff' argument of ?results
[2] see 'independentFiltering' argument of ?results

**RNA-seq Naive treated vs untreated**

out of 27456 with nonzero total read count
adjusted p-value < 0.05
LFC > 0 (up)       : 1255, 4.6%
LFC < 0 (down)     : 1701, 6.2%
outliers [1]       : 0, 0%
low counts [2]     : 10646, 39%
(mean count < 7)
[1] see 'cooksCutoff' argument of ?results
[2] see 'independentFiltering' argument of ?results

**RNA-seq EpiLC treated vs untreated**

out of 27456 with nonzero total read count
adjusted p-value < 0.05
LFC > 0 (up)       : 1015, 3.7%
LFC < 0 (down)     : 1198, 4.4%
outliers [1]       : 0, 0%
low counts [2]     : 12775, 47%
(mean count < 16)
[1] see 'cooksCutoff' argument of ?results
[2] see 'independentFiltering' argument of ?results

**GSEA: genes upregulated upon EpiLC induction±IAA against induction genes.**

| ID | setSize | enrichmentScore | NES | pvalue | p.adjust | qvalue | rank |
|---|---|---|---|---|---|---|---|
| Up | 972 | -0.5129332 | -2.181569 | 1e-10 | 1e-10 | NA | 3312 |

**GSEA: genes upregulated upon EpiLC induction±IAA against induction genes.**

| ID | setSize | enrichmentScore | NES | pvalue | p.adjust | qvalue | rank |
|---|---|---|---|---|---|---|---|
| Down | 1174 | 0.4596925 | 2.077525 | 1e-10 | 1e-10 | NA | 2891 |

**Comparison of lists:**

**Upregulated in EpiLC±IAA vs E6.5 male Het vs KO**

Fisher's Exact Test for Count Data

```
data:  matrix_data_up_RNA
p-value < 2.2e-16
alternative hypothesis: true odds ratio is not equal to 1
95 percent confidence interval:
 4.582795 7.345873
sample estimates:
odds ratio
   5.82226
```

**Downregulated in EpiLC±IAA vs E6.5 male Het vs KO**

Fisher's Exact Test for Count Data

```
data:  matrix_data_down_RNA
p-value < 2.2e-16
alternative hypothesis: true odds ratio is not equal to 1
95 percent confidence interval:
 4.765920 7.659175
sample estimates:
odds ratio
  6.059935
```

**Upregulated in Naive±IAA vs E6.5 male Het vs KO:**

Fisher's Exact Test for Count Data

```
data:  matrix_data_up_RNA_naive
p-value < 2.2e-16
alternative hypothesis: true odds ratio is not equal to 1
95 percent confidence interval:
 3.339051 5.375264
sample estimates:
```

odds ratio
4.25251

**Downregulated in Naive±IAA vs E6.5 male Het vs KO**

Fisher's Exact Test for Count Data

data: matrix_data_down_RNA_naive
p-value < 2.2e-16
alternative hypothesis: true odds ratio is not equal to 1
95 percent confidence interval:
3.513185 5.564747
sample estimates:
odds ratio
4.433634

**Upregulated in EpiLC±IAA (RNA-seq) vs FA±IAA (TTchem-seq)**

Fisher's Exact Test for Count Data

data: matrix_data_up_RNA
p-value < 2.2e-16
alternative hypothesis: true odds ratio is not equal to 1
95 percent confidence interval:
4.486911 5.829176
sample estimates:
odds ratio
5.11266

**Downregulated in EpiLC±IAA (RNA-seq) vs FA±IAA (TTchem-seq)**

Fisher's Exact Test for Count Data

data: matrix_data_down_RNA
p-value = 3.298e-11
alternative hypothesis: true odds ratio is not equal to 1
95 percent confidence interval:
1.445325 1.948665
sample estimates:
odds ratio
1.680618

**TBX3 Short inversion experiments**

Replicate 1

Kruskal-Wallis rank sum test

data: values by group

Kruskal-Wallis chi-squared = 13332, df = 2, p-value < 2.2e-16

Dunn Test

| .y. | group1 | group2 | n1 | n2 | statistic | p | p.adj |
|---|---|---|---|---|---|---|---|
| values | Empty | Wildtype | 5000 | 5000 | -115.4662 | 0 | 0 |
| values | Empty | Y298F | 5000 | 5000 | -57.7331 | 0 | 0 |
| values | Wildtype | Y298F | 5000 | 5000 | 57.7331 | 0 | 0 |

Replicate 2

Kruskal-Wallis rank sum test

data:  values by group
Kruskal-Wallis chi-squared = 13332, df = 2, p-value < 2.2e-16

Dunn Test

| .y. | group1 | group2 | n1 | n2 | statistic | p | p.adj |
|---|---|---|---|---|---|---|---|
| values | Empty | Wildtype | 5000 | 5000 | -115.4662 | 0 | 0 |
| values | Empty | Y298F | 5000 | 5000 | -57.7331 | 0 | 0 |
| values | Wildtype | Y298F | 5000 | 5000 | 57.7331 | 0 | 0 |
